# Retinoic acid fluctuation activates an uneven, direction-dependent network-wide robustness response in early embryogenesis

**DOI:** 10.1101/2020.07.15.203794

**Authors:** Madhur Parihar, Liat Bendelac-Kapon, Michal Gur, Tali Abbou, Abha Belorkar, Sirisha Achanta, Keren Kinberg, Rajanikanth Vadigepalli, Abraham Fainsod

## Abstract

Robustness is a feature of regulatory pathways to ensure signal consistency in light of environmental changes or genetic polymorphisms. The retinoic acid (RA) pathway, is a central developmental and tissue homeostasis regulatory signal, strongly dependent on nutritional sources of retinoids and affected by environmental chemicals. This pathway is characterized by multiple proteins or enzymes capable of performing each step and their integration into a self-regulating network. We studied RA network robustness by transient physiological RA signaling disturbances followed by kinetic transcriptomic analysis of the recovery during embryogenesis. The RA metabolic network was identified as the main regulated module to achieve signaling robustness using an unbiased pattern analysis. We describe the network-wide responses to RA signal manipulation and found the feedback autoregulation to be sensitive to the direction of the RA perturbation: RA knockdown exhibited an upper response limit, whereas RA addition had a minimal feedback-activation threshold. Surprisingly, our robustness response analysis suggests that the RA metabolic network regulation exhibits a multi-objective optimization, known as Pareto optimization, characterized by trade-offs between competing functionalities. We observe that efficient robustness to increasing RA is accompanied by worsening robustness to reduced RA levels and vice versa. This direction-dependent trade-off in the network-wide feedback response, results in an uneven robustness capacity of the RA network during early embryogenesis, likely a significant contributor to the manifestation of developmental defects.

## INTRODUCTION

Robustness is an important feature of regulatory pathways to ensure signal consistency in light of environmental changes or genetic polymorphisms. Retinoic acid (RA), a central regulatory signaling pathway active during embryogenesis and adult tissue homeostasis (Metzler and Sandell, 2016; le Maire and Bourguet, 2014), is synthesized from vitamin A (retinol, ROL) or other retinoids or carotenoids obtained from the diet (Kedishvili, 2016; Ghyselinck and Duester, 2019; Blaner et al., 2016; Fainsod and Kot-Leibovich, 2018). During embryogenesis, excessive or reduced RA signaling induces a wide array of embryonic malformations (Durston et al., 1989; Kessel and Gruss, 1991; Papalopulu et al., 1991; Marshall et al., 1992; Corcoran et al., 2002; Hollemann et al., 1998; Fainsod et al., 2020; Sive et al., 1990). Human disorders like vitamin A deficiency syndrome, DiGeorge syndrome, Fetal Alcohol Spectrum Disorder, Congenital Heart Disease, neural tube defects, and multiple types of cancer have been linked to reduced RA signaling (Hartomo et al., 2015; Kim et al., 2005; See et al., 2008; Urbizu et al., 2013; Pangilinan et al., 2014; Fainsod et al., 2020). RA levels are tightly regulated throughout life, controlling quantitative, spatiotemporal expression and activity of the RA metabolic network (Ghyselinck and Duester, 2019; Kedishvili, 2016; Blaner, 2019). The regulatory mechanisms that confer this tight regulation under constant environmental challenges, i.e., robustness, as well as conditions under which such mechanisms fail, are yet to be fully understood.

RA biosynthesis involves two sequential oxidation steps: first, mainly, multiple alcohol dehydrogenases (ADH) or short-chain dehydrogenase/reductases (SDR) can oxidize ROL to retinaldehyde (RAL), followed by a subfamily of aldehyde dehydrogenases, retinaldehyde dehydrogenases (RALDH) that catalyze the oxidation of RAL to RA (Kedishvili, 2016; Metzler and Sandell, 2016; Ghyselinck and Duester, 2019). ROL and RAL can also be produced from retinyl esters or ß-carotene from food sources (Blaner, 2019). RA availability is further regulated by several ROL, RAL, and RA binding proteins (Napoli, 2017). In gastrula vertebrate embryos, RA signaling is triggered by the initial expression of ALDH1A2 (RALDH2) completing the biosynthesis of RA (Niederreither et al., 1999; Begemann et al., 2001; Grandel et al., 2002; Chen et al., 2001; Shabtai et al., 2018). Substrate availability for the RALDH enzymes is controlled by multiple RAL reductases (Adams et al., 2014; Billings et al., 2013; Feng et al., 2010; Porté et al., 2013; Shabtai and Fainsod, 2018). Besides the temporal and spatial regulation of this wide array of RA metabolic network components, other RA network components, including the RA receptor families (RAR and RXR) and retinoid-binding proteins (Cui et al., 2003; Lohnes et al., 1995; Janesick et al., 2015) are under tight regulation. Then, RA signaling levels are controlled through its regulated biosynthesis and degradation, as well as restricted spatiotemporal expression of a network of metabolic and gene-regulatory components, which together are expected to provide robustness to RA signaling. The RA signal is sensitive to the maternal nutritional status and environmental exposure to chemicals (ethanol and others) (Shabtai et al., 2018; Paganelli et al., 2010), requiring the necessity to adapt the RA biosynthetic and signaling network to nutritional changes and environmental insults. This adaptation and maintenance of normal signaling levels under changing conditions is termed robustness (Nijhout et al., 2019; Eldar et al., 2004).

Over the years, multiple reports have supported the self-regulation of relatively few RA metabolic and signaling network components by RA itself. Among the network component genes shown to be responsive to RA levels, is *aldh1a2* whose expression has been shown to be down-regulated by increasing RA, in somite stage zebrafish embryos, neurula stage *Xenopus* embryos, and adult human epidermal cultures, or down-regulated or insensitive in mouse 8.5 dpc embryos, among others (Dobbs-McAuliffe et al., 2004; Chen et al., 2001; Niederreither et al., 1997; Pavez Loriè et al., 2009; Moss et al., 1998). A commonly studied counterpart of ALDH1A2 is the RA hydroxylase, CYP26A1, whose expression is commonly adjacent to the site of RA production (Moss et al., 1998; Hollemann et al., 1998). Up-regulation of the *cyp26a1* gene has been described in late gastrula *Xenopus* and zebrafish embryos, cranial up-regulation and caudal down-regulation in 8.5 and 9.5 dpc mouse embryos, human hepatocytes, and others (Moss et al., 1998; Hollemann et al., 1998; Fujii et al., 1997; Dobbs-McAuliffe et al., 2004; Topletz et al., 2015). Another important enzyme pair in RA metabolism commonly compared are DHRS3 and RDH10 which preferentially reduce RAL to ROL or oxidize ROL to RAL, respectively. Up-regulation of *dhrs3* was observed in late gastrula *Xenopus* embryos, while *rdh10* showed no change in chick and was down-regulated in *Xenopus,* both in neurula stage embryos (Kam et al., 2013; Reijntjes et al., 2010; Strate et al., 2009). Few studies have studied the effects of RA on the expression of other enzymes in the RA metabolic network. While these observations support an RA-dependent self-regulatory network, the multitude of experimental systems, range of developmental stages, and RA concentrations employed hamper the detailed understanding of the self-regulation of the RA network.

Many of the studies showing RA regulation of components of the RA metabolic network relied mainly on non-physiological and continued exposures to excess RA, and fewer studies employed a reduction of RA. These studies, although performed in different experimental systems, anatomical regions, developmental stages, and even under different experimental conditions, have led to insights on the self-regulation of the RA metabolic and signaling network. These studies provided initial understanding of the RA signaling robustness. However, the extent to which fluctuations in RA lead to network-wide feedback responses has not been investigated in-depth, including the limits of robustness achieved via feedback regulation. The common use of supraphysiological levels of RA to test the response of the embryo is largely due to the common observation that embryonic malformations are mild in response to fluctuations in RA within the physiological range (Sive et al., 1990; Durston et al., 1989; Kessel, 1992). RA knockdown studies taking advantage of inhibitors, inverse agonists, or enzymatic degradation (Janesick et al., 2014, 2013; Kot-Leibovich and Fainsod, 2009; Hollemann et al., 1998), revealed that the developmental malformations observed were usually milder than expected (Hollemann et al., 1998; Koide et al., 2001; Sharpe and Goldstone, 1997; Blumberg et al., 1997; Shabtai et al., 2018; Janesick et al., 2014). These observations suggest two possibilities to explain the mechanisms that can yield robustness to fluctuations in RA levels: insensitivity - the physiologically relevant fluctuations are too small to trigger a significant feedback response; or, active self-regulation of the RA metabolic network specifically aimed at overcoming the fluctuations and returning the RA levels to normal. To discriminate between these two alternative hypotheses, an unbiased RA network-wide analysis has to be performed to conclusively compare these scenarios. The promiscuity of RA network components for each step raises the possibility of partially overlapping robustness responses across biological replicates, and would suggest multiple alternative regulatory and enzymatic scenarios capable of returning the RA signal to normal developmental levels. The scarcity of studies performing a network-wide analysis means that the potential compositional complexity of the RA feedback response has not been truly explored.

In the present study, we have taken an integrated dynamic transcriptomics approach to address several unresolved questions regarding the RA signaling robustness mechanisms during early embryogenesis. We designed a pulse-chase, transient RA perturbation protocol to explicitly study the kinetics of the robustness response using an unbiased, transcriptomic-based (RNAseq), RA network-wide analysis of the robustness response to increased versus decreased RA levels. The composition of the RA metabolic network during gastrulation, and the individual and network-wide responsiveness to physiological RA manipulations were studied. We tested the limits of this robustness in both directions of RA perturbations and examined the clutch-to-clutch variations in terms of deviations from the normal response kinetics. Lastly, we combined the results from multiple clutches and different experimental techniques to propose a regulatory scheme that may underlie the robustness response.

## MATERIALS AND METHODS

### Embryo culture

*Xenopus laevis* frogs were purchased from Xenopus I or NASCO (Dexter, MI or Fort Atkinson, WI). Experiments were performed after approval and under the supervision of the Institutional Animal Care and Use Committee (IACUC) of the Hebrew University (Ethics approval no. MD-17-15281-3). Embryos were obtained by *in vitro* fertilization, incubated in 0.1% MBSH and staged according to Nieuwkoop and Faber (1967).

### Embryo Treatments

*all-trans* Retinoic acid (RA), Dimethyl sulfoxide (DMSO), and 4-Diethylaminobenzaldehyde (DEAB), were purchased from Sigma-Aldrich (St. Louis, Missouri). Stock solutions of RA, and DEAB, were prepared in DMSO. Two-hour treatments of 10 nM RA, or 50 µM DEAB, were initiated during late blastula (st. 9.5) and terminated at early gastrula (st. 10.25) by three changes of 0.1% MBSH and further incubation in fresh 0.1% MBSH for the desired time.

### Expression analysis

For each sample, 5-10 staged embryos were collected and stored at −80°C. RNA purification was performed using the Bio-Rad Aurum Total RNA Mini Kit (according to the manufacturer’s instructions). RNA samples were used for cDNA synthesis using the Bio-Rad iScriptTM Reverse Transcription Supermix for RT-qPCR kit (according to the manufacturer’s instructions). Quantitative real-time RT-PCR (qPCR) was performed using the Bio-Rad CFX384 thermal cycler and the iTaq Universal SYBR Green Supermix (Bio-Rad). All samples were processed in triplicate and analyzed as described previously (Livak and Schmittgen, 2001). All experiments were repeated with six different embryo batches. qPCR primers used are listed in (Table 1).

**Table 1.**
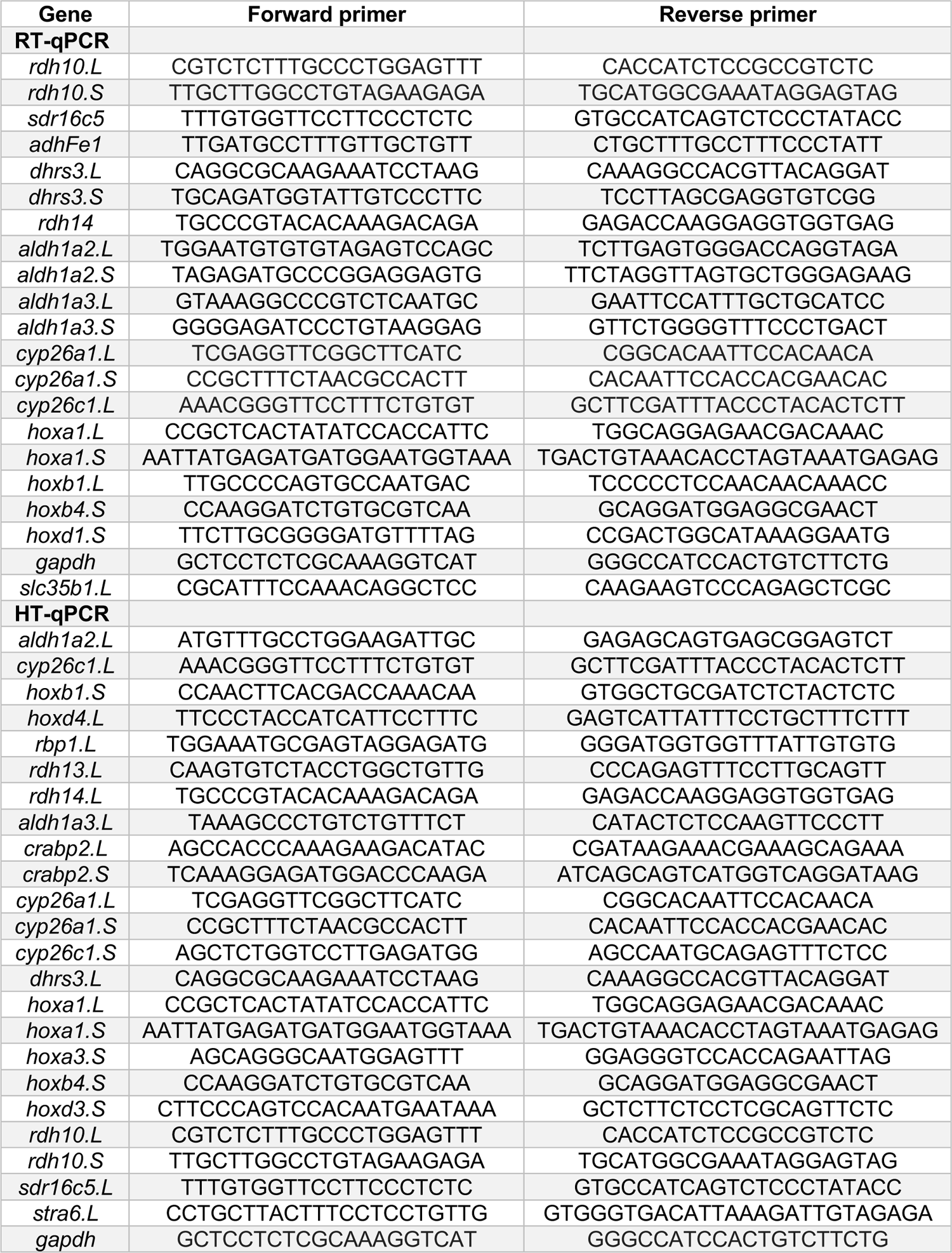
List of PCR primers corresponding to the *hox* genes and RA metabolic network for RT-qPCR and HT-qPCR analysis.

For spatial expression analysis of *hoxb1* and *hoxb4* we performed whole-mount *in situ* hybridization as previously described (Epstein et al., 1997). Probes for *in situ* hybridization used were: *hoxb1*, clone CX19/21 (Godsave et al., 1994) cut with EcoRI, transcribed with SP6 polymerase; *hoxb4*, clone E16 (Gont and De Robertis, personal communication) cut with BamHI, transcribed with T7 polymerase.

### RNASeq data analysis

Sequencing was performed at the Thomas Jefferson University Genomics Core using Illumina HiSeq 4000. Reads were mapped to the genome using the Xenopus laevis 9.1 genome with STAR alignment and a modified BLAST and FASTQC in the NGS pipeline (STAR average mapped length: 142.34). Annotation of the mapped sequences using *verse* identified 31535 genes. Raw counts were further filtered for non-zero variance across all samples resulting in 31440 scaffold IDs. We analyzed the time series transcriptomic data set for differentially expressed genes using a two-way ANOVA approach (time points: 0 (wash), 1.5, 3, 4.5 hours; treatments: RA, DEAB, control; n=6 biological replicates).

### High throughput qPCR

cDNA samples were directly processed for reverse transcriptase reaction using SuperScript VILO Master Mix (Thermo Fisher Scientific, Waltham, MA), followed by real-time PCR for targeted amplification and detection using the Evagreen intercalated dye-based approach to detect the PCR-amplified product. Intron-spanning PCR primers were designed for every assay using Primer3 and BLAST for 24 genes from Retinoic Acid metabolism and target pathway (Table 1). The standard BioMark protocol was used to preamplify cDNA samples for 22 cycles using TaqMan PreAmp Master Mix as per the manufacturer’s protocol (Applied Biosystems, Foster City, CA, USA). qPCR reactions were performed using 96.96 BioMark Dynamic Arrays (Fluidigm, South San Francisco, CA, USA) enabling quantitative measurement of multiple mRNAs and samples under identical reaction conditions. Each run consisted of 30 amplification cycles (15 s at 95°C, 5 s at 70°C, 60 s at 60°C). Ct values were calculated by the Real-Time PCR Analysis Software (Fluidigm). Samples were run in triplicate for the 24 genes. The primers in the first pre-amplification group selectively bind for the L homoeologues of the genes. In the second pre-amplification group, primers were selected for the S homoeologues only. Third pre-amplification group binds to all 24 genes. In order to remove the technical variability caused due to L and S gene homoeologues, Ct value for each gene in a sample was selected as the median value of the three pre-amplification runs. Relative gene expression was determined by the ΔΔCt method. *gapdh* was used as a housekeeping gene for normalization of the data.

### Data normalization and annotation

Data analysis on raw gene counts was performed using the R statistical analysis tool version 3.6.0 on a 64-bit Windows platform. For the RNA-seq data, the raw gene counts were first converted into log2-transformed values using the “regularized log (rlog)” transformation from the DESeq2 package, which minimizes differences between samples for rows with small counts (Love et al., 2014). The gene expression data was then normalized across samples against the experimental variation and batch effects using the COMBAT method in R using a non-parametric adjustment (Johnson et al., 2007). Following batch correction, the gene list was filtered for a minimum expression threshold to remove genes with normalized counts less than 5 across all 72 samples. The expression data for the remaining genes was normalized using quantile normalization. RNA-Seq transcript/array probes IDs were transformed to Official Gene Symbol using merged list from 3 sources: the *Xenopus laevis* scaffold-gene mapping, DAVID Bioinformatics Resource 6.8 (Huang et al., 2009) or AnnotationData package “org.Xl.eg.db” maintained by Bioconductor (Carlson, 2017). The original scaffold IDs were retained along with the Official Gene Symbols for cross-reference purposes.

### Differential gene expression analysis

The normalized data was analyzed using an Empirical Bayes Statistical (eBayes) model that considered the following two variables and their interactions: (1) Treatment (Control, RA, DEAB) and (2) Time post treatment-washout (t = 0, 1.5, 3, 4.5h) with n = 6 biological replicates. Differentially expressed genes were identified based on statistically significant effects of Time, Treatment or an interaction between these two factors. P-values were adjusted for multiple testing using topTable from *limma* (Ritchie et al., 2015) package in R (q < 0.05). The significane-filtered differential gene experssion data was used in an established Principal Component Analysis (PCA) approach using the *prcomp* function implemented in R. The samples were annotated based on a combination of treatment and time, yielding 12 distinct sample groups. For each of the selected PCs, expression of 100 top positively-loaded and 100 top negatively-loaded genes was visualized as a heat map.

### Dynamic pattern analysis and COMPACT analysis

First, for each time point, for each of the treatment conditions, the gene expression data for all six clutches (A, B, C, D, E, F) was averaged. Within treatment groups, RA, DEAB, control, the gene expression data at time points t = 1.5, 3, 4.5h was normalized by subtracting the corresponding ‘t=0h’ group. This average differential gene expression data was then discretized to three levels (+1, 0, −1) based on a fold-change threshold [±2 (up, no or down-regulation)]. Within the three treatment groups, this discretization yielded a dynamic response pattern vector for each gene, encoded by one of 27 (3 levels^3 time-points) possible ordered sets. Counts of genes in each treatment group that follow each of the 27 * 27 (=729) possibilities were compared. Functional enrichment analysis was performed for geneset in various dynamic pattern vectors in the Control conditions, using functions *enrichGO* and *simplify* from the R package “clusterProfiler” (Yu et al., 2012).

### COMPACT analysis of RA and DEAB after normalizing to Control

First, for each time point, for each of the treatment conditions, the gene expression data for all six clutches (A, B, C, D, E, F) was averaged. Within treatment groups, RA and DEAB, the gene expression data were normalized to Control at each time point by subtracting the expression of the Control group at the corresponding time point. This Control-normalized differential gene expression data for both treatment groups was then discretized to three levels (+1, 0, −1) based on a fold-change threshold [+log2(1.3) (up, no or downregulation)]. This discretization yielded a dynamic response pattern vector for each gene, encoded by one of 81(3 levels^4 time-points) possible ordered sets. Subsequently, RA and DEAB groups were compared to count the number of genes corresponding to each of the 81 * 81 (=6561) possibilities; to create an 81 × 81 matrix representing the comparative dynamic response pattern counts (COMPACT) (Kuttippurathu et al., 2016). For a given COMPACT matrix of comparative conditions RA(vs. Control) and DEAB (vs. Control), the element at the ith row and jth column of the matrix contains the number of genes that show an ‘i’th pattern in DEAB and ‘j’th pattern in RA. For a coarse-grained version of the detailed 81×81 COMPACT, pattern-vector counts for each treatment group were further aggregated based on the first time-point, yielding 9 groups of pattern vectors per treatment group. The pair of treatment groups (RA and DEAB) was then compared to count the number of genes corresponding to each of the 9 * 9 (=81) possibilities.

### RA network map and visualization

A schematic representation for the position and function of genes involved in RA biosynthesis, metabolism, translocation, and transcription during gastrula stages was derived by data mining. The gastrula expression of each gene was confirmed by qPCR. For each time point, for each treatment, expression value for each gene was mapped to the corresponding label in the schematics using a color scale.

Gene correlation and clustering analysis - Gene expression data was analyzed for both with and without Control-normalization using Weighted Gene Coexpression Network Analysis (WGCNA) (Langfelder and Horvath, 2008) to identify modules of genes with highly correlated differential expression. We used a soft threshold value of 9 (8 for Control normalized data) to identify the initial gene coexpression modules, followed by a dissimilarity threshold of 0.25 to merge the initial modules into the final set of gene coexpression modules. Identified modules were further correlated with the traits (batch, time, treatment).

### Robustness score calculation and principal curve-based trajectory analysis

RNA-seq expression data of (clutches: A,B,C,D,E,F) and additional normalized qPCR expression data (clutches: G,H,I,J,K,L) were independently genewise Z-score transformed. Transformed data was then combined to result into 144 samples (12 clutches * 3 treatments * 4 time points). We further selected two subsets of genes: 1) “RA metabolism genes” [*aldh1a2.L, aldh1a3.L, crabp2.L, crabp2.S, cyp26a1.L, cyp26a1.S, cyp26c1.L, cyp26c1.S, dhrs3.L, rbp1.L, rdh10.L, rdh10.S, rdh13.L, rdh14.L, sdr16c5.L, stra6.L*]. This set represents the feedback regulatory mechanism utilized in response to the RA levels perturbations. 2) “HOX genes” [*hoxa1.L, hoxa1.S, hoxa3.S, hoxb1.S, hoxb4.S, hoxd4.L*] representing the phenotypic outcome from the treatment. For each gene set, the PCA scores for the combined data was first calculated using R function *prcomp* and then the scores from the first three principal components (PC1, PC2 and PC3) are used to learn a 3-dimensional principal curve using the function *principal_curve* from R package princurve. Two sets of points are specified for the ‘start’ parameter for this function, which determines the origin and direction of the curve: (1) The centroid of the 0h-Control samples from all twelve clutches, and (2) centroid of all the remaining 132 samples. For each geneset (RA metabolism genes and HOX genes), for each clutch (A-L), for each treatment (RA or DEAB), a “net absolute expression shift” can be calculated as the sum of distances along the principal curve, of the treatment samples from the corresponding control samples:

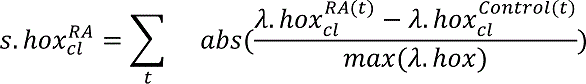

where, *λ. hox_cl_^RA(t)^* is the arc-distance from the beginning of the curve, for the “*RA treatment samples*” at time point “*t*”, for clutch “*cl*” for the principal curve learned for the geneset “*Hox genes*” (Returned as the parameter “*lambda*” from the principal curve function). The expression shift calculation allows ranking and sorting clutches based on ‘HOX shift’ (*S. hox_cl_^treatment^*) and ‘RA Network shift’ (*S. metabolism_cl_^treatment^*). A higher HOX shift for a clutch for a treatment indicates that the clutch is more robust to that particular treatment/perturbation. Furthermore, clutches were mapped onto the conceptual map across the spectrum of “HOX shift” and “RA Network shift” divided into hypothetical quadrants of Effective or Ineffective regulation, or Efficient or Inefficient regulation.

## RESULTS

### Transient physiological perturbations to study RA signaling robustness

To directly investigate the robustness of RA signaling, we designed a pulse-chase RA manipulation protocol to transiently perturb this signaling pathway beyond its normal, and developmentally changing levels, and study the feedback regulatory changes involved in restoring its normal signaling levels (Fig. 1A). The network-wide changes of the transient RA disturbances and restoration of normal signaling levels were monitored by kinetic transcriptomic analysis (RNAseq). Parameters such as developmental stages and timing between samples were empirically optimized (not shown). This experimental approach expands on reports describing RA-driven regulation of its own metabolic and signaling network components. To reduce RA levels below the normal content in early embryos, we partially inhibited the oxidation of RAL to RA with the RALDH inhibitor 4-diethylaminobenzaldehyde (DEAB) (Shabtai et al., 2016). To increase RA levels for robustness analysis in a physiologically-relevant manipulation, we treated embryos with 10 nM *all*-trans RA which is below the reported physiological RA content of early/mid gastrula *Xenopus laevis* embryos (about 100-150 nM all-*trans* RA) (Durston et al., 1989; Kraft et al., 1995; Creech Kraft et al., 1994, 1995; Kraft et al., 1994; Schuh et al., 1993; Chen et al., 1994).

**Figure 1.**
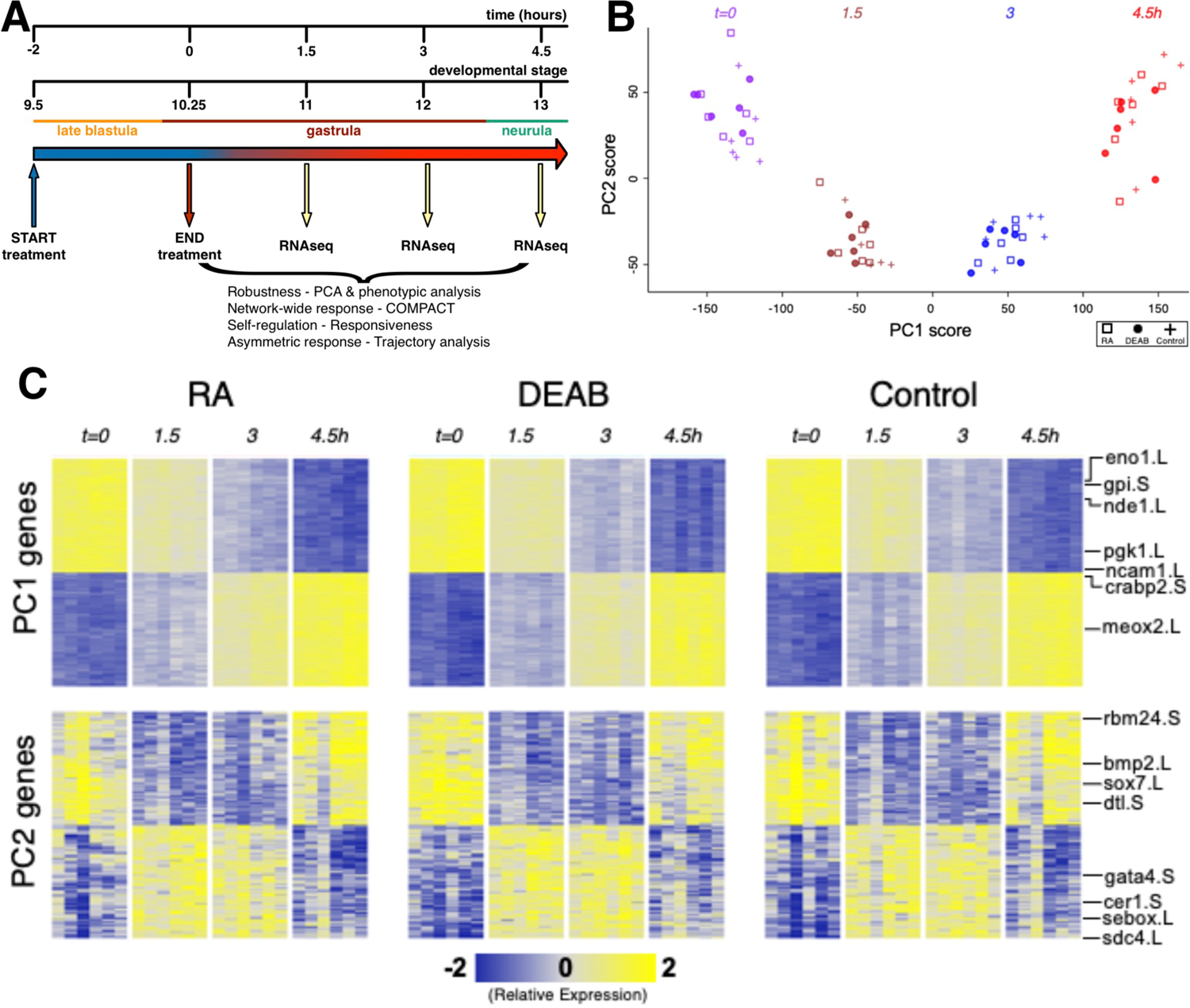
Efficient RA signaling robustness following transient physiological manipulation of RA levels. **(A)** Schematic depiction of the experimental design to directly challenge and study the robustness of RA signaling during embryogenesis. Timing and developmental stages are shown. **(B)** Principal Components 1 and 2 (PC1, PC2) of all six biological replicates studied by time series RNAseq. **(C)** Heatmap of gene expression of the top-100 positive and top-100 negative loadings corresponding to PC1 and PC2. A subset of the genes is highlighted based on the relevance to early developmental processes.

Transient pharmacological RA manipulations of early *Xenopus* embryos were performed by treating with RA (10 nM) or DEAB (50 µM) for 2 hours from late blastula (st. 9.5; t=-2h) to early gastrula (st. 10.25; t=0h), washed, and samples collected every 1.5 hours (t=1.5, 3, 4.5h) after treatment termination (Fig. 1A). For each biological repeat, all treatments, controls, and time points were collected from a single clutch (eggs) of a single female. Treatment efficiency and quality of the RNA samples was first monitored by quantitative RT-PCR (qPCR) for changes in *hox* gene expression as an internal readout of the changes in RA signal levels. Only those biological repeats where both treatments exhibited the expected upregulation (RA) or downregulation (DEAB) of *hox* gene expression were selected for RNAseq analysis.

Principal Component Analysis (PCA) of the gene expression variation showed separation along the first principal component (PC1) which corresponds to transcriptomic changes of the normal progression through gastrulation (Fig. 1B,C). RA-manipulated samples clustered with the control samples of the same developmental stage suggesting a robust maintenance of transcriptome composition. Normal transcriptomic changes as a result of progression through embryogenesis appear to be the dominant variable distinguishing the samples irrespective of the RA manipulation (Fig. 1B,C). The second component (PC2) separated the sample groups at intermediate time points (t=1.5 and t=3h) from those at t=0 and 4.5h, indicating a transient differential expression shift as the next most dominant pattern in the data (Fig. 1B,C). These results show that overall, the dynamic transcriptome is maintained along the normal developmental trajectory despite the perturbations to RA levels.

The effects of RA and DEAB on the transcriptome were not readily apparent in the first seven principal components. PC8 showed some separation of the DEAB group from the RA and Control groups (Supplemental Fig. 1A,C), whereas PC10 showed some separation of the RA treatment group from Control and DEAB treatments (Supplemental Fig. 1B,D). The top-ranked genes along PC8 and PC10 showed distinct dynamic expression patterns across the treatments, whereas top-ranked genes along PC1 and PC2 largely corresponded to in-common dynamic changes over time (Fig. 1B; Supplemental Fig. 1C). The overall magnitude of induced changes in gene expression in response to RA or DEAB appears to be less than the normal transcriptome changes occurring during early developmental stages. These observations suggest that RA signaling robustness dampens the gene expression changes otherwise induced by abnormal RA levels in agreement with a robust system that functions to limit the gene expression changes in the majority of the transcriptome.

### Retinoic acid signaling exhibits efficient robustness

Further support for the robustness of RA signaling was obtained by phenotypic and molecular analysis of embryos treated for RA knockdown. *Xenopus* embryos treated with the RALDH inhibitor, DEAB (50 µM), or overexpressing CYP26A1 to render RA inactive (Hollemann et al., 1998; Yelin et al., 2005; Topletz et al., 2015) developed with unexpectedly mild defects for such an important signaling pathway (Figs. 2B,C,E). The induction of severe phenotypes by RA loss-of-function required the combined RA knockdown with both treatments targeting the metabolism at different steps (Fig. 2D,E) supporting the observed robustness of RA signaling. Analysis of changes in *hoxa1* or *dhrs3* expression supported the conclusion that the individual RA-knockdown treatments result in moderate gene expression changes (Fig. 2F-H), and severe gene expression changes are observed when both treatments were combined (Fig. 2F-H). These results further support the notion that RA robustness needs to be overcome to elicit strong gene expression and phenotypic changes.

**Figure 2.**
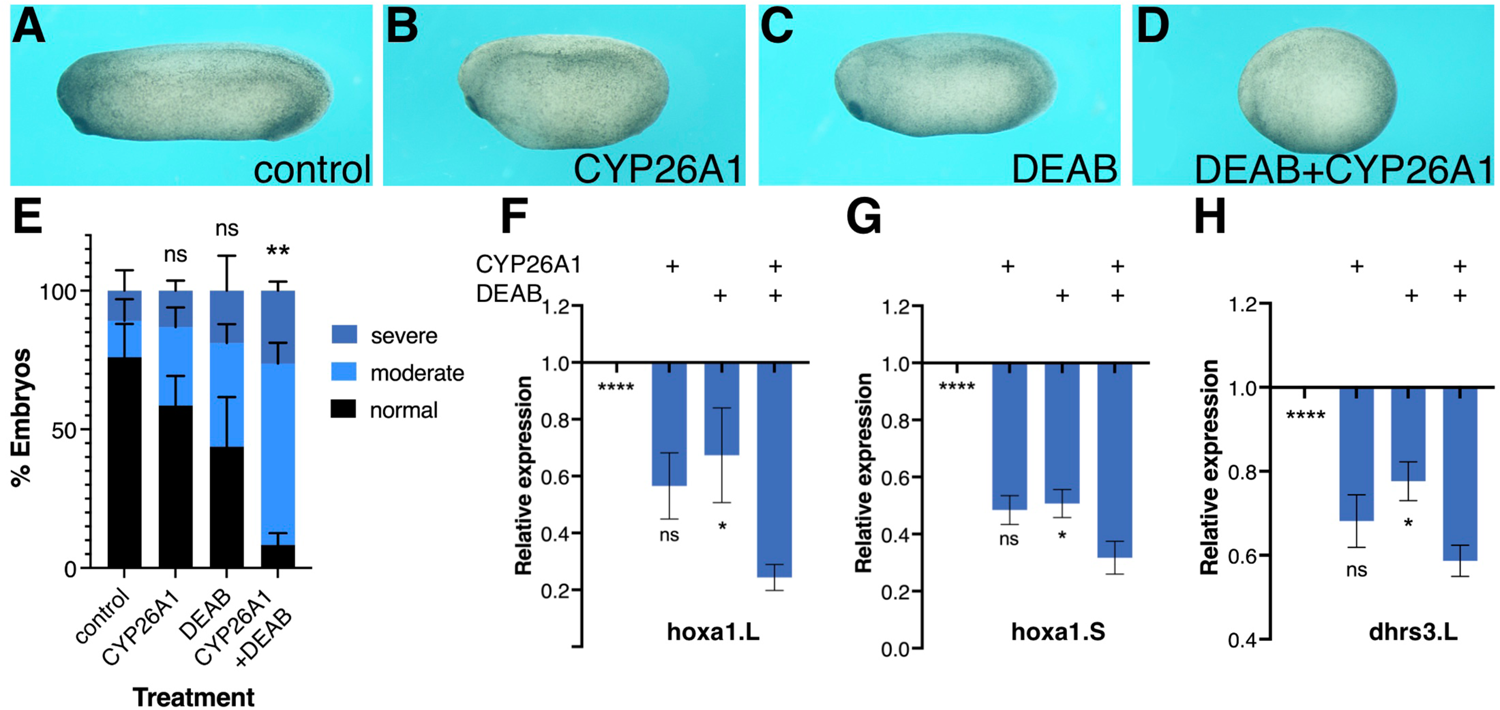
Phenotypic and molecular robustness of the retinoic acid metabolic pathway. Retinoic acid levels were reduced in *Xenopus laevis* embryos by inhibition of the RALDH activity with DEAB or by CYP26A1 overexpression to render retinoic acid inactive. **(A)** Control embryo at st. 27. **(B)** Embryo injected with capped RNA (0.8 ng) encoding the CYP26A1 enzyme. **(C)** Embryo treated with DEAB (50 µM) from st. 8.5 until st. 27. **(D)** Embryo treated with DEAB and injected with *cyp26a1* mRNA. **(E)** Distribution of the developmental defect severity induced by the retinoic acid manipulations. Embryos were scored for the induction of moderate or severe phenotypes, or normal looking. **(F-H)** Gene expression changes as a result of retinoic acid manipulation. qPCR analysis of *hoxa1.L* **(F)**, *hoxa1.S* **(G)**, and *dhrs3.L* **(H)** relative expression levels as a result of the individual or combined retinoic acid manipulations. Statistical significance (Student’s t-test) was calculated compared to the combined treatment group. *, p<0.05; **, p<0.01; ****, p<0.0001; ns, not significant.

To begin exploring the limits of the robustness response to increased RA levels we treated embryos with increasing RA concentrations in the physiological range and above (1 nM - 1 µM). Treated embryos were analyzed for morphological malformations by early tailbud stages (st. 32-33) and revealed that the RA treatments induced developmental malformations whose severity increases in correlation to the amount of RA added (Fig. 3A,B). While exposure to 10 nM RA resulted in very mild malformations involving mainly the anterior head (forebrain) region, 50 nM RA induced a clear and relatively strong phenotype, and 100 nM RA resulted in severe defects, both affecting the head (microcephaly) and trunk (anterior-posterior axis) regions (Fig. 3A,B). In similarly treated embryos, we analyzed the expression changes of genes encoding RA metabolic enzymes and RA targets (*hox* genes) during early (st. 10.25) and late gastrula (st.12) (Nieuwkoop and Faber, 1967). Genes positively regulated by RA at both stages, *hoxb1*, *cyp26a1*, and *dhrs3* exhibited dose-dependent responses (Fig. 3C-E). The genes, *aldh1a2* and *rdh10,* were down-regulated by most RA doses (Fig. 3F,H), while *aldh1a3* (*raldh3*) was upregulated at early gastrula and was repressed late gastrula (Fig. 3G). Importantly, in many instances, treatments above 10 nM RA resulted in stronger changes in gene expression or the transition to more severe malformations suggesting an upper limit to the robustness response. We also studied the effects of graded increase in RA levels on the spatial expression patterns of *hoxb1* and *hoxb4* (Supplemental Fig. S2). We performed a visual estimation of the expression level changes compared to control embryos, or compared to the previous (lower) RA concentration. As determined by qPCR for *hoxb1* (Fig. 3C), for both genes we observed the expected RA concentration-dependent *in situ* hybridization signal intensity increase (Supplemental Fig. S2). In parallel, we observed a widening of the expression domain of both genes along the anterior-posterior axis (Supplemental Fig. S2A-C,E-G).

**Figure 3.**
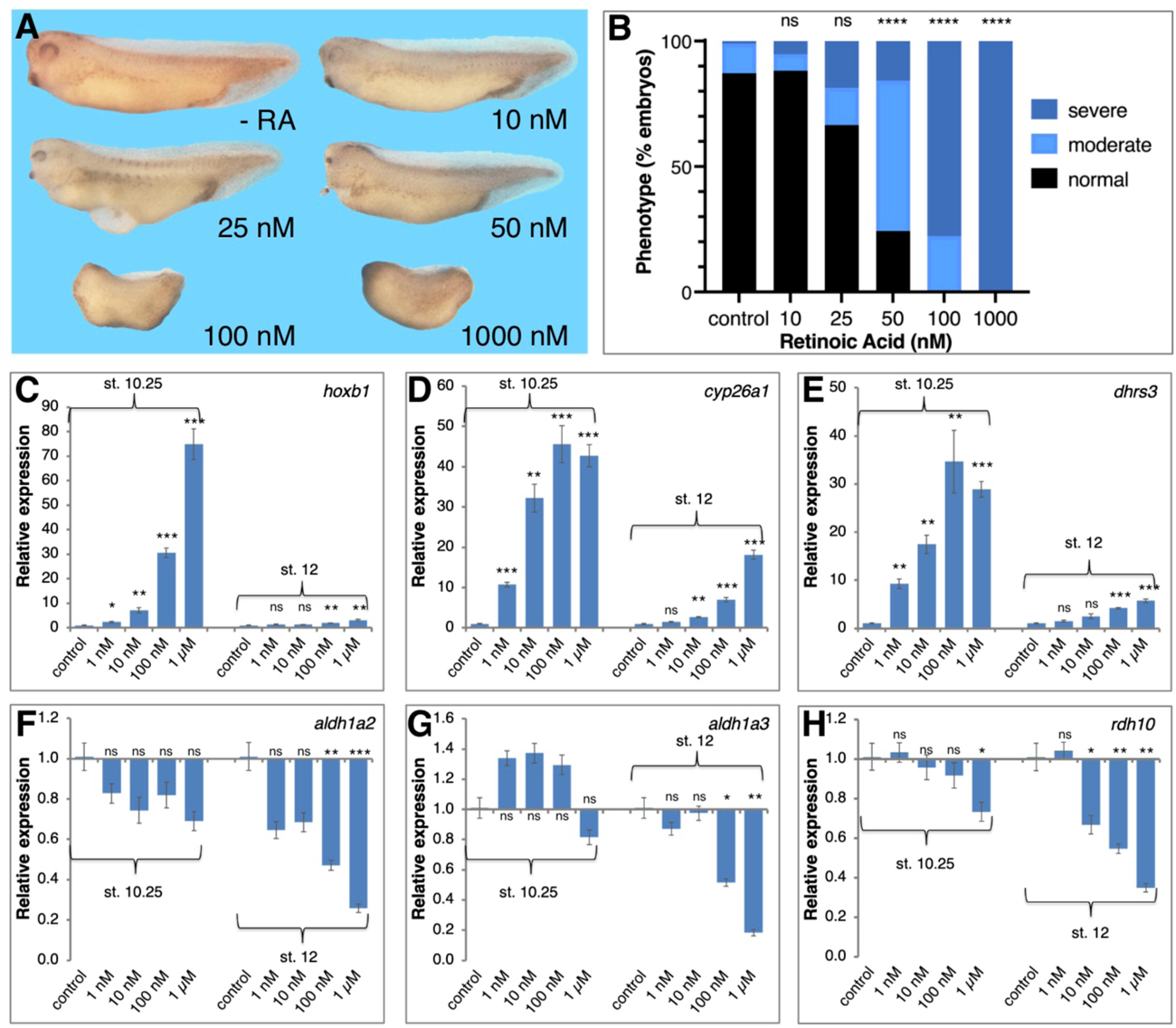
Transient physiological manipulation of RA levels for robustness analysis. **(A)** Embryos were treated with increasing concentrations of all-*trans* RA from 1 nM to 1 µM. Treatments were initiated at st. 8.5 and RNA samples were collected at tailbud stages (st. 32-33) for phenotypic analysis. **(B)** Distribution of the severity developmental defects induced by the retinoic acid manipulations. Embryos treated were scored for the induction of moderate or severe phenotypes, or normal looking. **(C-H)** RA treated embryos were collected at early (st. 10.25) and late (st. 12) gastrula for analysis of the changes in expression level. The response of RA metabolic and target genes was studied by qPCR. **(C)** *hoxb1* **(D)** *cyp26a1* **(E)** *dhrs3* **(F)** *aldh1a2* **(G**) *aldh1a3* **(H)** *rdh10*. Statistical significance (Student’s t-test) was calculated compared to the control group. *, p<0.05; **, p<0.01; ***, p<0.001; ns, not significant.

Further support for the robustness of RA signaling came from the in-depth analysis of the kinetic transcriptomic changes. We analyzed the averaged data for all biological repeats using an unbiased dynamic pattern analysis approach along all possible discretized patterns of gene expression (Kuttippurathu et al., 2016). Following normalization to t=0h, up- or down-regulated genes beyond the significance threshold at each time point were scored as either +1 or −1, respectively or 0 for no significant change (Supplemental Fig. S3A). Such an exhaustive approach allows us to enumerate the dominant as well as subtle patterns in the data overcoming limitations of conventional cluster analysis that could miss or mask the smaller groups of genes with distinct expression profiles over time.

Expression of a total of 4693 genes significantly changed more than 2-fold over time (compared to t0; multiple testing corrected q-value < 0.05), grouped in only 10 patterns out of the possible 27 (3*3*3) dynamic patterns for any of the experimental groups (RA, DEAB, Control) (Supplemental Fig. S3A). Only 5 patterns were common enough to include 100 or more genes in any experimental group, and they corresponded to up- or down-regulation at later time points (3h and 4.5h) suggesting a return to normal expression levels after the initial RA-dependent disruption (Supplemental Fig. S3A). There were no significant differences in the gene counts between RA or DEAB and control groups (two-tailed Z test, p > 0.05). Gene Ontology enrichment of the control group highlighted development, cell cycle, morphogenesis, RA biosynthesis, gastrulation, and others, as expected (Supplemental Fig. S3B, Supplemental Table S1). At this level of analysis, we could not observe distinctive dynamic gene expression patterns that occur only in response to the RA or DEAB treatments. Venn diagram analysis revealed limited (about 20-30%) treatment-specific specific gene expression changes that do not overlap between treatment and control samples for each pattern. For instance, in the subset of genes that showed late downregulation (pattern 0,0,-1; Supplemental Fig. S3C) only 216 genes in DEAB or 187 genes in RA, did not overlap with the control sample (1175 genes) showing that only about 18% of the genes changed as a result of the treatments beyond the normal developmental changes. The expression changes relative to the end of the RA perturbation (t=0h) did not vary substantially (Supplementary Figure 3), consistent with the observed robustness.

### RA network-wide responses to restore normal signaling levels

We exhaustively compared the RA and DEAB treatments adapting our unbiased COMPACT approach for analyzing time-series differential transcriptomic profiles across multiple experimental conditions (Kuttippurathu et al., 2016). For statistically significant changing genes within each treatment group (two-way ANOVA; q<0.05), we constructed discrete patterns based on differential gene expression between treatment (RA or DEAB) and control at each time point (81 theoretically possible patterns). Also here, only 32 distinct patterns with at least one gene in either comparison were observed (Fig. 4A). The effect threshold, 1.3-fold average difference, was chosen lower than 2-fold, as the differential expression analysis revealed that the RA/DEAB perturbations were leading to a smaller magnitude of changes at each time point, as compared to the larger changes occurring normally over time (Fig. 1B, Supplementary Fig. S3), providing additional evidence of the robustness. Intersecting the two distributions yielded a matrix where each element corresponds to a distinct pair of patterns corresponding to the gene-specific response to increased (RA vs. control) and decreased (DEAB vs. control) RA signaling (Fig. 4B).

**Figure 4.**
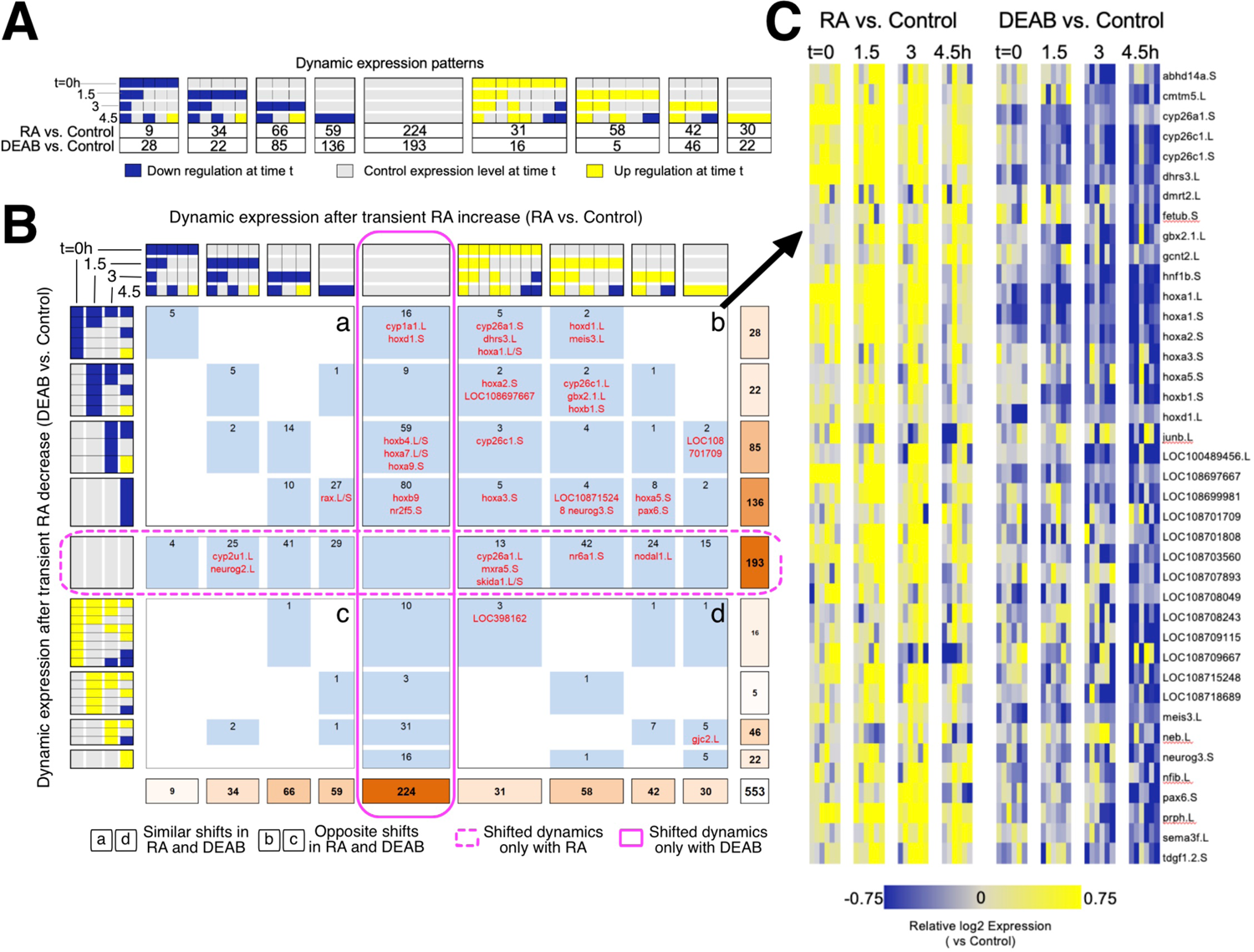
Comparative pattern analysis to uncover genes responding to opposing RA manipulations. **(A)** Genes were grouped into discretized expression patterns based on up-regulation (yellow), down-regulation (blue), or no change (grey) compared to the control sample at the same time point. The numbers below the patterns indicate the number of genes that show the corresponding expression pattern in RA vs. control or DEAB vs. control. Only 32 out of 81 theoretically possible (four time points, 3*3*3*3=81) dynamic patterns were exhibited by at least one gene and are included in the figure. **(B)** COMPACT matrix comparing gene expression changes due to RA and DEAB relative to control. The subset of 32 x 32 patterns with non-zero number of genes in either perturbation group are shown. The gene counts were grouped within related patterns based on the time of initial up- or down-regulation. Supplemental Data 1 contains a version of the COMPACT matrix shown with the gene identifiers corresponding to the counts. The different quadrants are labeled a to d. **(C)** Dynamic expression patterns of genes showing opposite changes in response to RA and DEAB treatments (quadrant b).

Surprisingly, the COMPACT matrix (Fig. 4B, Supplemental Data 1) shows that the majority of the genes sensitive to RA manipulation exhibited sensitivity to only one direction of the RA perturbation. A total of 193 genes only responded to the addition of RA (middle row), whereas 224 genes were only responsive to RA knockdown (middle column). Of the genes responsive to both RA and DEAB perturbations, 88 genes responded in the same direction to both treatments (quadrants a and d; Fig. 4B). Only a set of 48 genes showed opposite transcriptional outcomes to RA increase versus RA decrease (quadrants b and c; Fig. 4B). Importantly, the set of genes that showed upregulation by exogenous RA addition and downregulation by DEAB (quadrant b; Fig 4B) was enriched for genes involved in RA metabolism (Fig. 4B,C).

Analysis of the RA signaling robustness revealed a complex, network-wide transcriptional response aimed at altering enzyme levels to redirect the metabolism of RA towards restoration of normal signaling levels. To better understand the network-wide response we proceeded to determine the components of the RA network expressed during early gastrula to specifically analyze them from the transcriptomic data. The RA network composition during early/mid gastrula was determined by data mining starting from a list of 27 possible RA metabolic and signaling network components. For 12 of the 27 genes, we found transcriptomic support in our data or additional RNAseq data sets (Xenbase.org) (Yanai et al., 2011; Savova et al., 2017; Karimi et al., 2018). All putative RA network components were validated by determining their temporal expression pattern by qPCR, analyzing both homeologues (L and S genes) when present in the *X. laevis* genome (Supplemental Fig. S4). The basic RA metabolic network analyzed includes during early/mid gastrula 12 genes out of which 4 are singletons and 8 are present as L or S homeologs (Fig. 5A) (Session et al., 2016).

**Figure 5.**
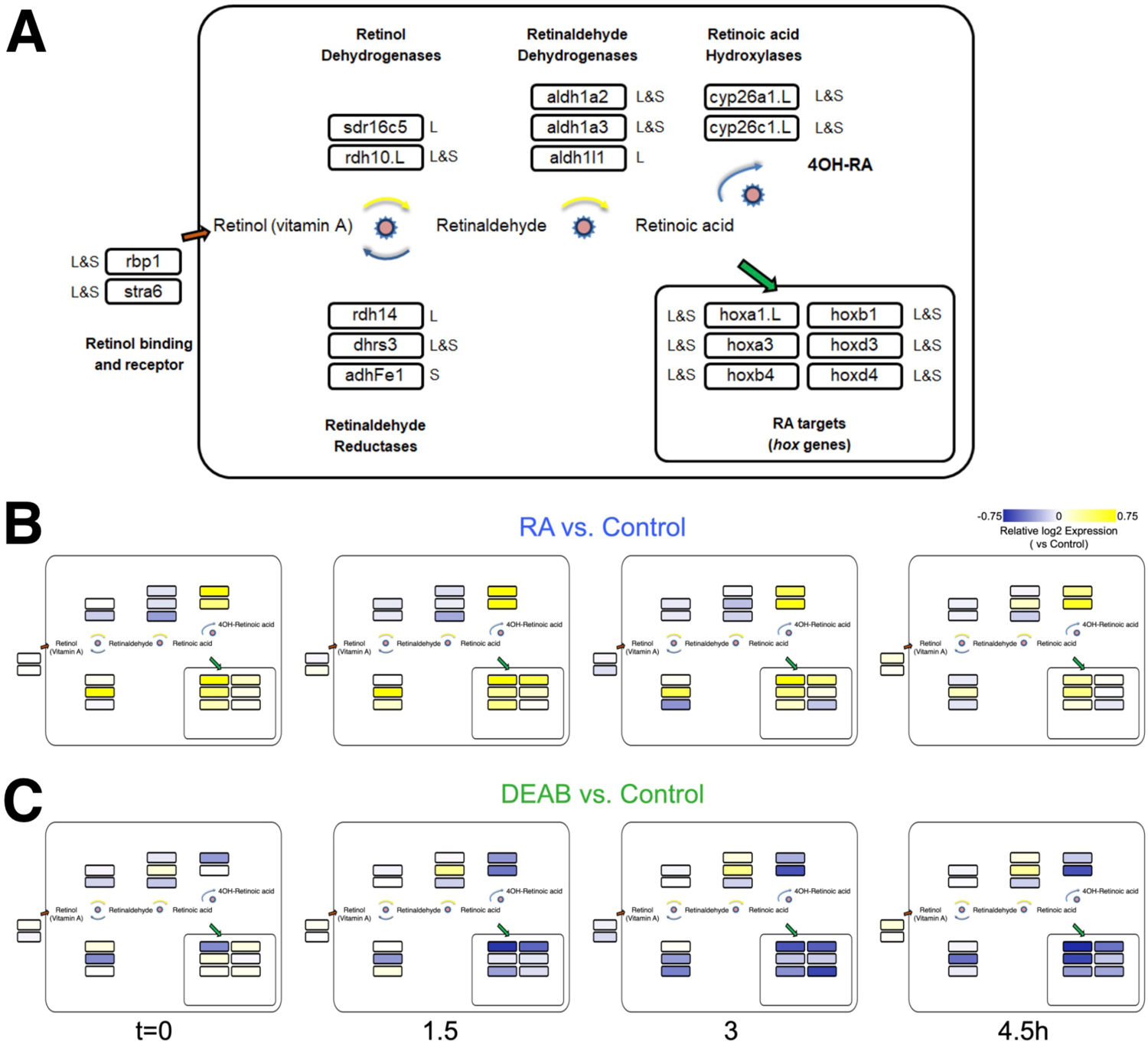
Network-wide response to transient RA manipulations. **(A)** RA metabolic network components and RA target genes analyzed during early/mid gastrula. The homeolog (L&S) or singleton status of the gene(s) in the genome is denoted outside the box with the gene name. The gene names shown include a .L suffix to mark in cases where only one of the homeologs changed significantly in the RNAseq and HT-qPCR data. **(B,C)** Mapping the differential expression data onto the RA network shown in panel **(D)** to highlight the differences in regulation of biosynthesis versus degradation of RA signal between RA and DEAB treatment groups.

We mapped the expression changes in response to RA perturbation (Fig. 4B,C) to the RA metabolic network (Fig. 5). Notably, based on their reported preferential enzymatic activity, not all genes exhibited their expected gene expression change (Fig. 5B,C). For instance, following transient RA addition, genes encoding enzymes with a preferential retinaldehyde reductase activity (*rdh14* and *adhFe1*) undergo downregulation, and not the expected upregulation (Fig. 5B). As an independent validation, we analyzed the differential expression time series data using another unbiased approach, Weighted Gene Correlation Network Analysis (WGCNA) (Langfelder and Horvath, 2008). A WGCNA gene expression module with the highest correlation with treatment contained a similar gene set as shown in Fig. 4C, supporting the COMPACT analysis (green module, Supplemental Table S2; Supplemental Data 2). Taken together, our results support a network-wide mechanism to maintain RA signaling levels to counter the effects of external perturbations and prevent teratogenic effects by a transcriptional feedback control system. Many of the feedback regulatory interactions uncovered require further characterization.

To cross-validate the observed kinetics of the RA robustness response, we performed the pulse-chase RA perturbations in other independent clutches of embryos (Fig. 1A) and analyzed the expression changes by qPCR (Supplemental Fig. S5). In RA-treated embryos, the two RA-regulated genes, *hoxb1* and *hoxb4,* were upregulated at the time of treatment washing (t=0h) (Supplemental Fig. S5A), and two hours into the recovery (t2), both RA target genes were back to control expression levels. We studied the expression of *dhrs3*, which preferentially reduces RAL to ROL to attenuate RA biosynthesis (Feng et al., 2010), *cyp26a1*, which targets RA for degradation, and *aldh1a2* that produces RA from RAL (Shabtai et al., 2016), to begin understanding the RA robustness response. Both negative RA regulators of RA biosynthesis, *dhrs3,* and *cyp26a1* were upregulated at t0 by increased RA and return to normal expression levels after a 4-hour recovery (Supplemental Fig. S5B). As expected, *aldh1a2* exhibited an RA-promoted weak downregulation at t0, returning to normal levels after 4 hours (Supplemental Fig. S5B). These results exemplify how the gene expression levels of RA metabolic enzymes are regulated to achieve normal RA levels as part of the robustness response. A complementary study performed by inhibiting RA biosynthesis with DEAB (Supplemental Fig. S5C,D) revealed that all genes studied exhibited fluctuations during the recovery period. The genes *hoxd1* and *hoxb1* reached almost normal levels at an earlier stage than the genes encoding RA metabolic components, again showing a concerted, network-wide, response of the RA metabolic network to restore normal developmental RA levels and *hox* gene expression (Supplemental Fig. S5C). Genes encoding anabolic enzymes, e.g., *aldh1a2*, were upregulated, and catabolic components, e.g., *cyp26a1*, were downregulated (Supplemental Fig. S5D), consistent with a network-wide response model of adjusting the pathway component expression (and activity) to restore normal RA signaling levels.

### Direction-dependent uneven RA robustness response efficiency

To better characterize the gene responses observed in the RNAseq study to RA manipulation, we performed a detailed titration of RA levels and determined the response of the main RA network genes. Multiple embryo clutches (n=5) were exposed to increasing concentrations of RA (1, 10, 100, and 1000 nM) or several concentrations of DEAB (1, 5, 10, 25, and 50 µM) to induce a gradual reduction in RA levels during mid gastrula (st. 11). Gene expression changes were studied by qPCR for both homeologues (Fig. 6A-F). The results show that genes encoding RA production limiting enzymes (Dhrs3) or enzymes promoting RA degradation (Cyp26a1) exhibit a robust, dose-dependent response to increased RA similar to well-characterized RA target genes such as *hoxa1* (Fig. 6A-C). In contrast, as a result of RA reduction, no clear response trend could be observed, but by 25 µM DEAB, expression of both, *dhrs3* and *cyp26a1* decreased by about 30% similar to *hoxa1* (Fig. 6A-C). For genes encoding enzymes active in RA production, the situation is more complex. *rdh10* exhibited a dose-dependent reduction trend with increasing RA concentrations (Fig. 6D). In contrast, reducing RA levels did not uncover a clear regulatory response. During mid gastrula, *aldh1a2* exhibited a mixed response to different levels of increased or decreased RA (Fig. 6E). Also, *aldh1a3* is down-regulated as a result of RA addition but no clear trend is observed and exhibits a mixed response to RA reduction (Fig. 6F). These results recapitulate in more detail the basic observations of the COMPACT (Fig. 4B) and the pattern response analysis (Supplemental Fig. S3A) where the majority of the genes responded to only one direction of RA perturbation.

**Figure 6.**
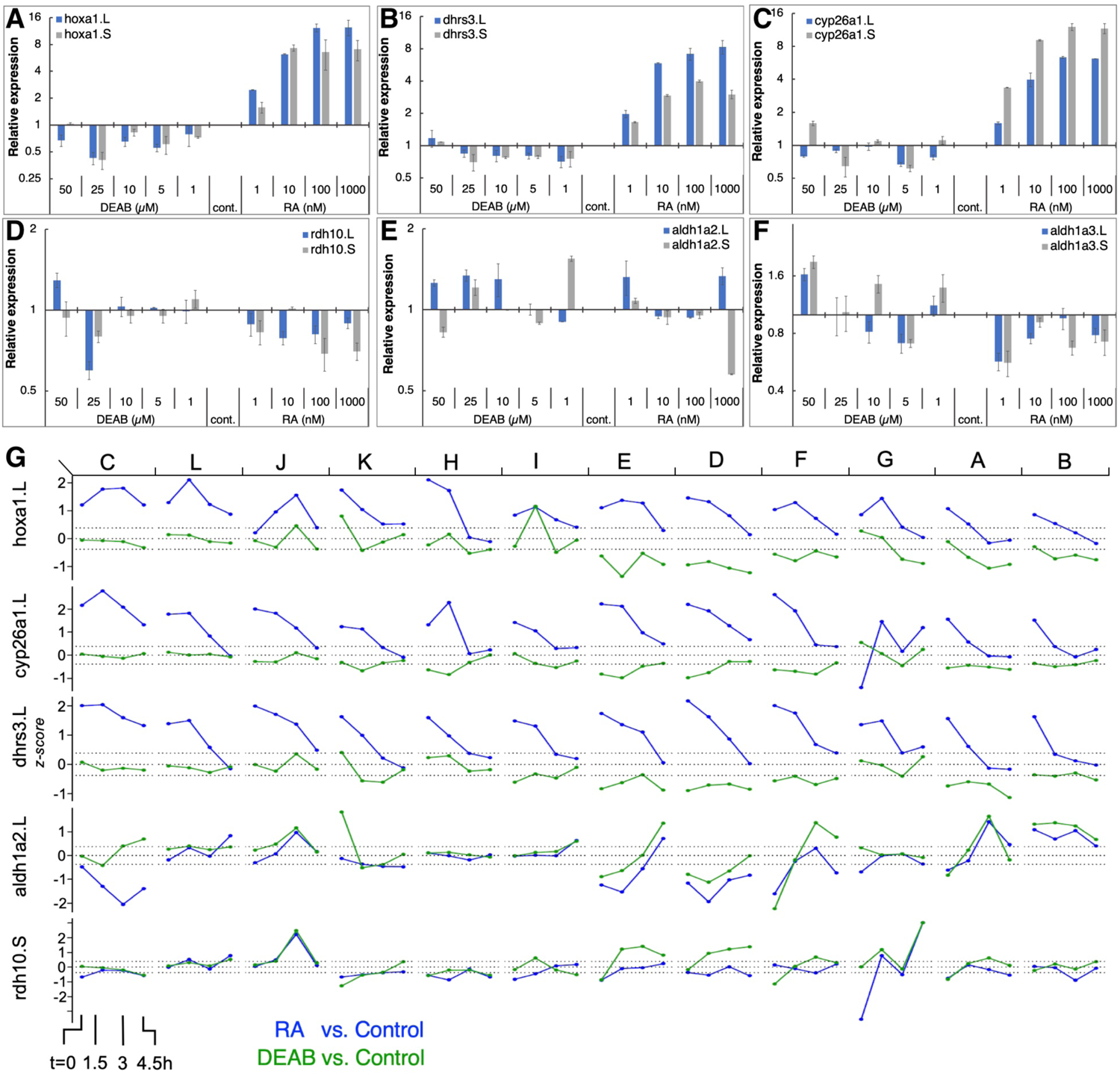
RA responsiveness and differential expression dynamics of select RA network genes and targets in different clutches. **(A-F)** Embryos were treated at midblastula (st.8.5) with increasing concentrations of RA (1-1000 nM) or DEAB 1-50 µM) to generate a gradient of RA responses. At mid gastrula (st. 11) the changes in RA network genes was determined by qPCR. **(A)** *hoxa1.L, hoxa1.S*; **(B)** *dhrs3.L, dhrs3.S*; **(C)** *cyp26a1.L, cyp26a1.S*; **(D)** *rdh10.L, rdh10.S*; **(E)** *aldh1a2.L, aldh1a2.S*; **(F)** *aldh1a3.L, aldh1a3.S.* **(G)** Dynamic expression patterns of multiple RA network genes across the different clutches. The data was combined from RNAseq (clutches A-F) and HT-qPCR (clutches G-L). The clutches are ordered left to right based on the earliest time at which *hoxa1* expression returned to the baseline levels.

The PCA analysis revealed that all samples (RA, DEAB, and control) from the same biological repeat (clutch) tended to cluster together, but the different clutches separated slightly from each other suggesting some clutch-dependent heterogeneity in gene expression (Fig. 1B; Supplemental Fig. S6). For this reason, we analyzed the transcriptional changes in the RA metabolic network within each clutch from the RNAseq data. In parallel, we generated an additional set of biological replicates (n=6) to further study the dynamic expression changes in RA metabolic network genes using high-throughput real-time PCR (HT-qPCR; Fluidigm Biomark; Supplemental Fig. S7) to validate the RNAseq results. These six additional biological repeats (labeled G-L) were treated with RA or DEAB following the same experimental design used for the RNAseq study (Fig. 1A). We observed a similar heterogeneity between clutches analyzed by HT-qPCR to the clutch-to-clutch variation in the RNAseq data (labeled A-F) (Supplemental Figs. S6 and S7).

We ordered the 12 clutches according to the earliest time point at which *hoxa1* returned to control levels (Fig. 6G and Supplemental Fig. S8) intermingling clutches of both studies. We observed high variability in the response to the RA manipulation among the individual clutches. Interestingly, clutches with significant *hoxa1* upregulation following RA addition showed limited downregulation in response to RA biosynthesis inhibition (clutches C, L, J, K, H, I in Fig. 6G). Clutches E, D, F, G, A, B (Fig. 6G) showed the opposite response, i.e. robust *hoxa1* change following RA knockdown, while RA increase induced a milder response. The RA suppressors, e.g., *cyp26a1* and *dhrs3*, showed significantly altered expression in clutches with larger deviations in *hoxa1* expression (C, L, J, K, H) and a strong and extended response to the addition of RA. In contrast, the differential expression of RA producers (e.g., *aldh1a2* and *rdh10*) after RA manipulation was relatively mild and did not fully align with the deviation in *hoxa1* expression (Fig. 6G). Such a heterogeneous correlation between the extent of feedback regulation and clutch-to-clutch variation in *hoxa1* expression was observed across the RA metabolic network (Supplementary Fig. S8).

### Uneven robustness in response to increased versus decreased RA signaling

To better understand the differences in the response of the different clutches to RA manipulation, we sought to quantitatively rank the robustness of each clutch in an integrated manner based on the kinetics of the expression shift of multiple *hox* genes and to analyze the feedback response of the RA metabolic network genes. We implemented a trajectory-based approach in which the samples of all biological repeats (clutches) at all time points were projected onto the first three principal components based on the expression of *hox* genes (*hoxa1*, *hoxa3*, *hoxb1*, *hoxb4*, *hoxd4*) or the RA metabolic network genes (Fig. 7A and Supplemental Fig. S9A). A principal curve was fit to the projected data, representing the trajectory in which the system evolves over time for all clutches combined. Each clutch was visualized separately along the principal trajectory, allowing us to compare the temporal evolution of deviations between RA manipulations and controls (Fig. 7A and Supplemental Fig. S9A). As a multi-gene measure of the robustness of each clutch, the net absolute distance between samples and controls was computed at all time points along the principal curve. The distance between treatments and controls decreased over time, with significant differences between clutches in the time taken to close the gap (Fig. 7B and Supplemental Fig. S9B). The wide range of clutch-to-clutch robustness variability assessed by analysis of multiple *hox* genes showed that clutches with low robustness to increased RA showed larger distances between samples and controls at any time point (Fig. 7A, clutch C), compared to robust clutches with decreasing distances (Fig. 7A, clutch E). In contrast, DEAB treatment resulted in an opposite robustness pattern (Fig. 7A), with clutch C showing the least deviation and clutch E the widest deviation (Fig. 7A bottom row and Fig. 6B). Clutch J exhibited intermediate robustness responses to both RA manipulations (Fig. 7A,B). The anti-correlated pattern of RA robustness suggests an uneven tradeoff in the gastrula embryo to counter changes in RA levels (Fig. 7C). The *hox* responses to altered RA levels revealed that all biological repeats aligned along a negative-slope diagonal and covered the whole range of responses (Fig. 7C). This result suggested the establishment of an RA robustness gradient among the clutches and a trade-off in the intrinsic capacity to counteract increased or reduced RA levels, i.e., uneven robustness.

**Figure 7.**
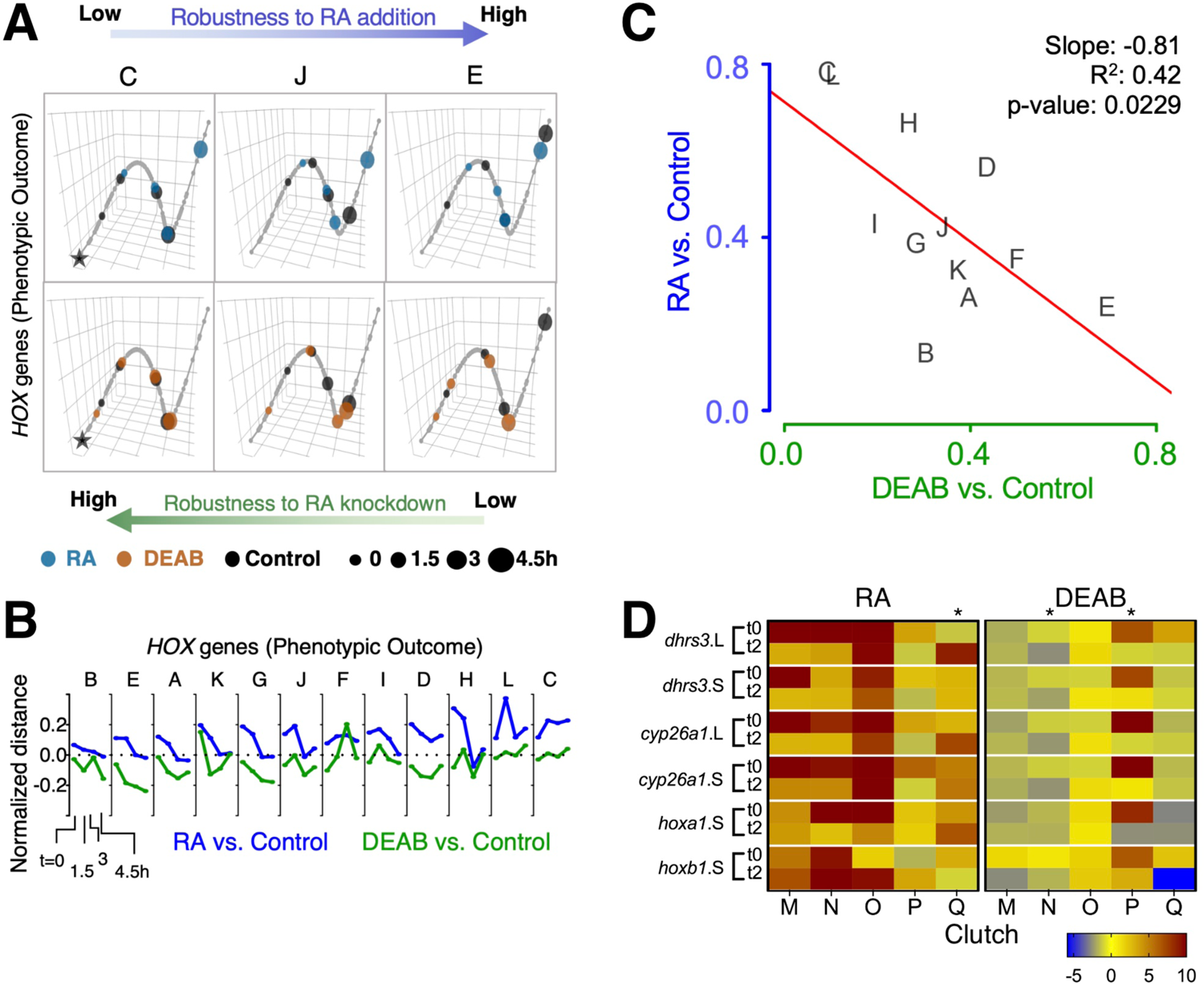
Trajectory analysis to compare the extent of individual clutch robustness based on *hox* expression. **(A)** 3-dimensional principal curve for the *hox* genes, showing projections of the sample points on the curve for **(A, top)** RA and **(A, bottom)** DEAB treatments. Principal curves for clutches C, J, and E are shown. The black star indicates the beginning of the curve for the distance measurement along the trajectory. Ranking of clutches is based on the net (absolute) normalized distance of treatment samples from the corresponding control sample for each time point. **(B)** Normalized expression shift profile calculated from the principal curve as the arc distance between the treatment and the corresponding control. Clutches are rank-ordered from lowest to highest net expression shift for *hox* genes in the RA group. Clutches A-F data from RNA-seq, clutches G-L data from HT-qPCR. **(C)** Distribution of clutches based on the trajectory determined robustness to RA and DEAB treatments relative to each other. The letters indicate the distinct clutches. **(D)** Heat map representation of gene expression changes of 5 additional clutches transiently treated with RA or DEAB. Samples were analyzed at the end of the pulse (t0) and 2 hours in the chase (t2). Asterisks denote clutches with abnormal responses.

A similar analysis of the genes encoding RA network components showed a scattered pattern among the biological repeats with no clear trend, but a characteristic pattern of reduced deviation over time in several clutches (Supplemental Figs. S9A,B). There was no significant correlation in the net shift in feedback regulatory action between RA addition versus reduction (Supplemental Fig. S9C). Importantly, the scattered clutch distribution depended on the direction of the RA manipulation. The range of clutch distribution for DEAB treatment suggests a response to very small RA reductions and an upper response limit (Supplemental Fig. S9C). In contrast, the clutch distribution in response to RA addition suggests that the response requires a minimal threshold change to be activated and it also exhibits an upper threshold. These observations suggest a high sensitivity to any amount of RA reduction but a lower sensitivity to small increases in RA levels.

Independent validation of these observations was performed by transiently treating multiple additional embryo clutches (n=5; M-Q) to disturb RA levels (RA or DEAB). Individual gene responses were analyzed by qPCR at the end of the treatment (t0) and 2 hours into the recovery from the treatment (t2). Following transient RA treatment, the expression changes of RA-responsive genes revealed that in the majority of cases we could observe the expected initial up-regulation and the subsequent relative down-regulation (Fig. 7D). As we observed previously in the RNAseq and HT-qPCR individual clutch analysis where some clutches (L, H, G) exhibited apparently reversed responses (Fig. 6G and Supplementary Fig. S8), also in this case, clutch Q exhibited a relative up-regulation during the recovery period (Fig. 7D). Following the transient reduction of RA levels (DEAB), we also observed clutches (N and P) that did not exhibit the characteristic up-regulation during the recovery period (Fig. 7D), but rather exhibited a relative down-regulation like clutch K (Fig. 6G and Supplementary Fig. S8). Importantly, the same clutches with abnormal responses (Q, N, P), exhibited normal responses to the complementary RA manipulation supporting the uneven response to increased versus decreased RA levels.

### Robustness variability impacts the efficiency-efficacy of the response to RA manipulation

Plotting the robustness response outcome (trajectory analysis) of the RA target genes (*hox* genes) (Fig. 6A) as a function of the change in the RA metabolic network required to achieve the recovery from the RA manipulation (Supplemental Fig. S9A) can provide insights on the effort invested by each clutch to achieve its own specific level of RA signaling robustness (Fig. 8A,B). Analysis of the response to increased RA levels showed some correlation between the RA network and *hox* shifts (Fig. 8A). Interestingly, increased RA prompted a strong feedback response in 9 out of 12 of the clutches, but only half of them achieved a high robustness outcome (Fig. 8A). Also, two clutches exhibited a highly efficient robustness response with a mild feedback response (Fig. 8A). In agreement with the proposed uneven robustness response, for reduced RA levels, the outcome (*hox* shift) as a function of the network shift revealed no clear correlation (Fig. 8B). While we observe *hox* responses, i.e. robustness outcomes, throughout most of the range, the network response exhibits an upper limit (Fig. 8B). This clutch response distribution suggests that although the RA network exhibits a mild feedback response for all the clutches, this is sufficient to achieve a sensitive, high robustness to reduced RA levels in at least 8 out of 12 clutches (Fig. 8B; lower left quadrant). The comparison of the robustness response (*hox*) and the effort to achieve it (network response) yielded an efficiency-efficacy matrix, within which clutches whose network response was low and the *hox* shift was also minimal were identified as establishing an efficient robustness response (Fig. 8C). At the other extreme, some clutches exhibited extensive shifts in the RA network response that were unable to minimize the *hox* shifts resulting in an ineffective robustness response (Fig. 8C). The different robustness responses of the clutches analyzed suggested that for each clutch, the robustness response is the result of a dynamic adaptation of the components of the network available. We examined whether neighboring clutches in the efficiency-efficacy matrix (Fig. 8C) employ similar strategies for feedback regulation. Interestingly, we found that the identity of the differentially regulated genes in the RA metabolic network can vary between clutches co-localized in the matrix (Figs. 8D,E). For example, in response to increased RA, clutches A, E and F showed similar differential regulation of *dhrs3*, *aldh1a2,* and *cyp26a1*, but diverged in *stra6*, *sdr16c5*, *adhfe1*, *rdh13*, *aldh1a3*, and *cyp26c1* (Figs. 8A,D). Similarly, in response to reduced RA, clutches A, B and D showed downregulation of *dhrs3* and *cyp26a1*, and upregulation of *rbp1*, suggesting feedback aimed at reducing enzymes active in RA reduction, and increasing import of retinol. However, there was high variability in the differential expression of *aldh1a2* and *aldh1a3* to increase RA production (Figs. 8B,E).

**Figure 8.**
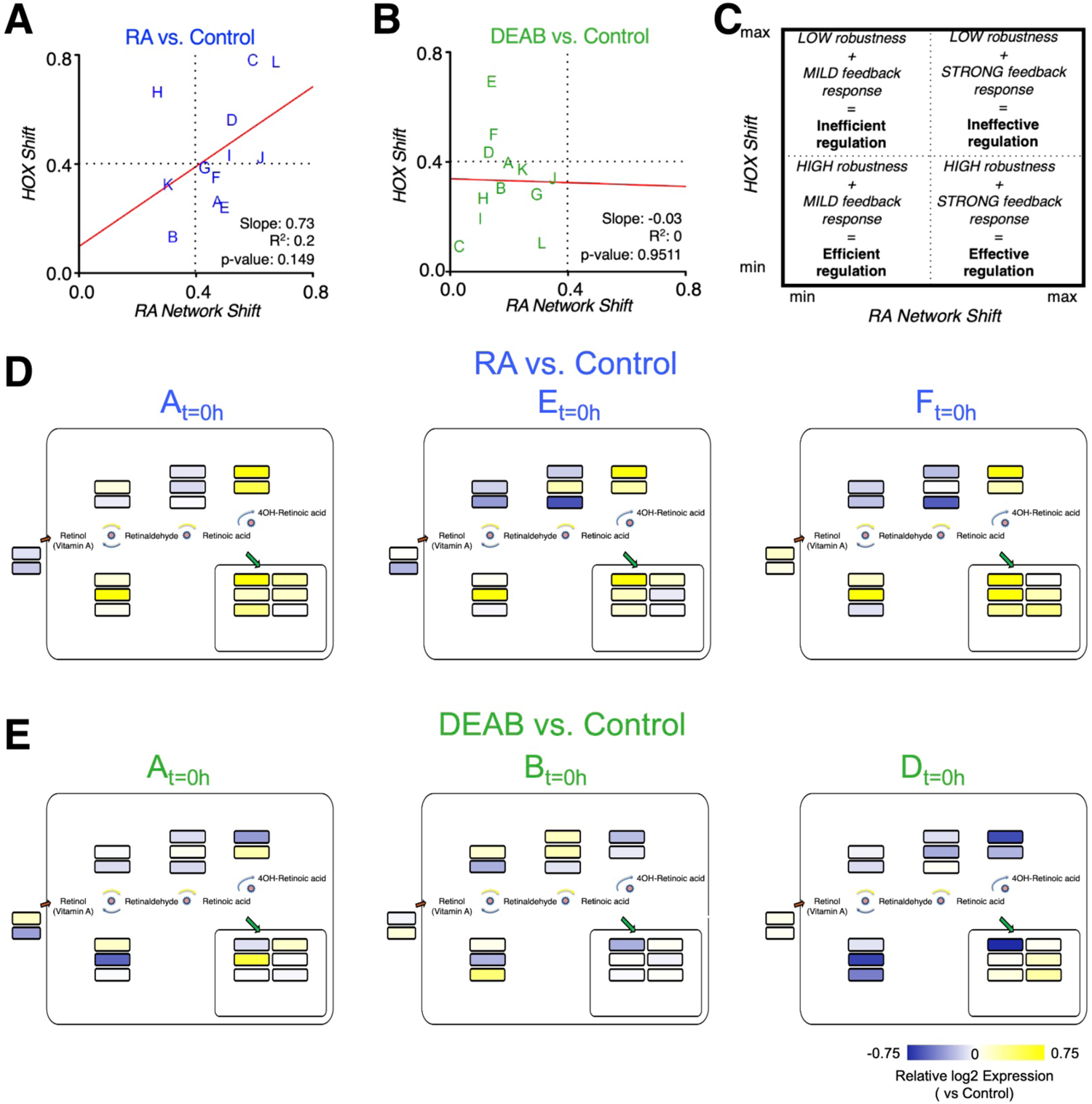
Differential robustness to direction of RA change is related to the effectiveness of feedback regulatory action. **(A,B)** Comparative distribution of clutches based on the trajectory determined robustness to RA **(A)** and DEAB **(B).** The change of the *hox* genes as a function of the change in the RA network genes is plotted for each RA manipulation. The letters indicate the distinct clutches. **(C)** Schematic diagram of the robustness efficiency-efficacy matrix in which different quadrants indicate the relative level of feedback observed and robustness achieved. **(D,E)** Mapping the differential expression data of clutches A, E, and F onto the RA network shown in Fig. 6D to highlight the heterogeneity of differential regulation of RA network components for clutches closely situated in the robustness efficiency matrix **(A,B).**

The observations of the uneven and only partially overlapping transcriptomic changes between clutches led us to begin exploring the alternative RA-regulated expression shifts of individual network components to achieve signaling robustness (Fig. 9). Based on the analysis of clutch-specific responses, we propose a regulatory scheme to help explain how different alternative strategies of changes to the RA network activity can lead to similar robustness levels. This decision tree exemplifies how each clutch follows a decision trajectory based on the direction of RA fluctuation and clutch-specific composition of RA network variants (Fig. 9). In this decision tree scheme, most robust RA network-wide adjustments in response to increase versus decrease in RA follow distinct non-overlapping trajectories. From our results on the gradient of clutch-wise robustness (Fig. 6C), it follows that each clutch is likely optimized to follow a trajectory in the decision tree with a trade-off for following other trajectories, such that high robustness to one direction of RA fluctuation may be achieved at the cost of low robustness to opposite direction of perturbation in RA levels, i.e., uneven robustness. In addition, clutches could also be optimized for following multiple trajectories with reduced robustness capacity in each scenario, yielding moderate robustness to either direction of RA fluctuation. Taken together, these results provide strong evidence that the early embryo is capable of robust maintenance of RA levels by mounting a range of network-wide feedback regulatory responses, which are constrained by an apparent trade-off that unevenly limits robustness for one of the directions of RA fluctuation.

**Figure 9.**
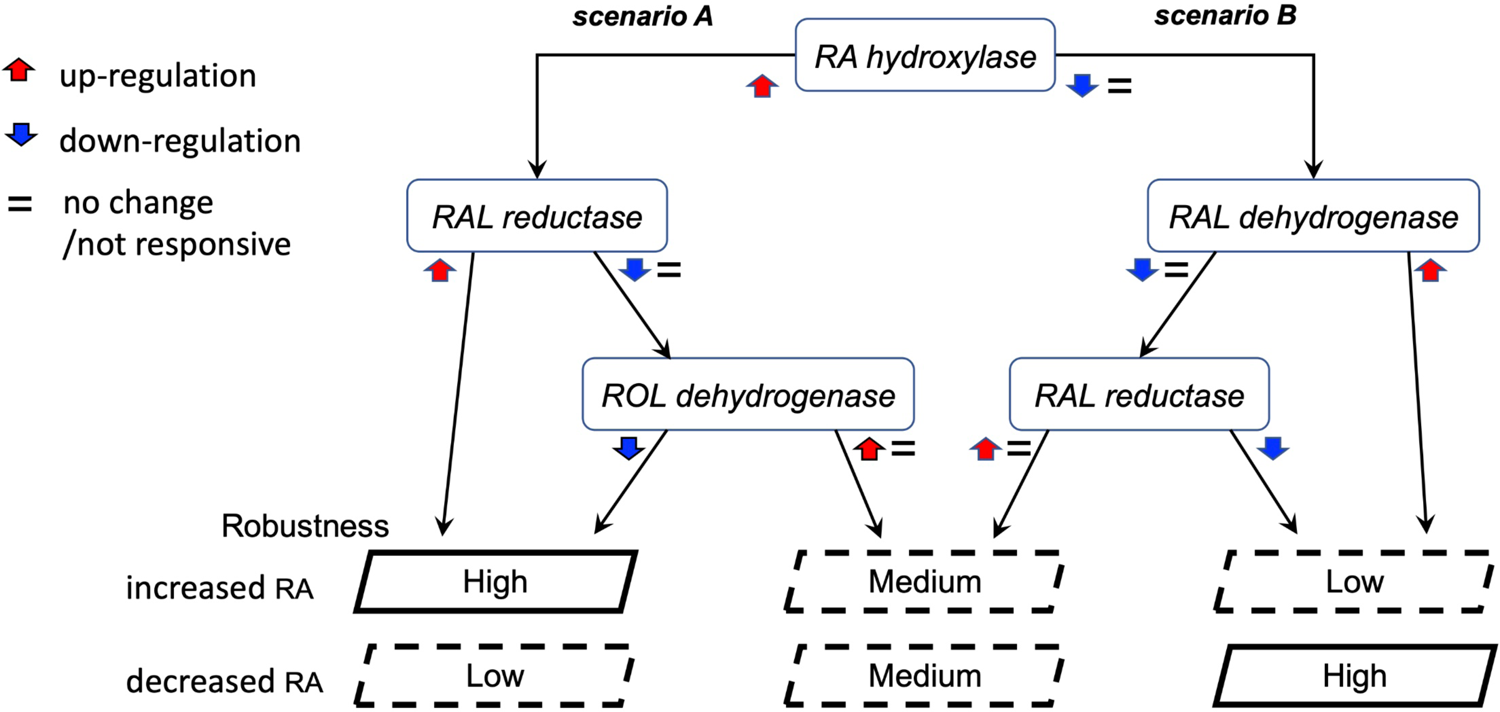
Proposed model of the alternative regulatory schemes to achieve RA robustness. Schematic representation of two alternative scenarios in the decision tree two achieve robustness. These two scenarios are representative of the multiple permutations possible to achieve similar robustness outcomes. Each clutch follows a decision trajectory based on the direction of RA fluctuation and clutch-specific composition of RA network variants. In this scheme, uneven robustness arises when the regulatory feedback response of a clutch is optimized for one scenario over another. Our results revealed such a trade-off over distinct objectives of counteracting RA increase versus decrease.

## DISCUSSION

### Robustness of the retinoic acid signaling pathway

Signaling pathway robustness is aimed at ensuring signaling consistency and reliability overcoming changing environmental conditions or genetic polymorphisms resulting in gene expression changes or protein activity variation. Commonly, strong developmental defects by RA manipulation in *Xenopus* embryos, and many other systems, are induced by treating with high RA concentrations in the 1 µM to 10 µM range (Sive et al., 1990; Taira et al., 1994). This is in contrast to the estimation that gastrula stage *Xenopus laevis* embryos contain RA levels around the 100 nM - 150 nM range (Durston et al., 1989; Kraft et al., 1995; Creech Kraft et al., 1994, 1995; Kraft et al., 1994; Schuh et al., 1993; Chen et al., 1994). To study the physiological robustness of RA signaling, we determined that by treating embryos with as little as 10 nM RA (about 10% of the estimated endogenous content) we could consistently induce measurable, even if less than dramatic, expression changes in RA-regulated genes. Our results and other studies have shown that embryo treatments with physiological RA concentrations exhibit relatively mild developmental malformations suggesting the activation of compensatory mechanisms to prevent abnormal gene expression and developmental malformations (Sive et al., 1990; Hollemann et al., 1998; Reijntjes et al., 2005). Similar mild defects were observed when we reduced RA levels by either blocking the biosynthesis (DEAB treatment) or by targeting this ligand for degradation (CYP26A1 overexpression), and these observations are supported by multiple loss-of-function studies describing mild developmental malformations induced by RA signaling reduction (Hollemann et al., 1998; Koide et al., 2001; Sharpe and Goldstone, 1997; Blumberg et al., 1997; Shabtai et al., 2018; Janesick et al., 2014). Therefore, the mild developmental defects induced by physiological increase or decrease in RA levels can be explained by a compensatory transcriptional response, i.e., robustness. An alternative explanation could be an intrinsic insensitivity of RA-mediated gene regulation to physiological fluctuations. Our results reject the latter explanation and support the former model by showing activation of a compensatory transcriptional robustness response. We show that one approach to overwhelm this RA signaling robustness is to interfere with the metabolic/signaling network at multiple steps (e.g., DEAB and CYP26A1 induction) to hamper its ability to efficiently elicit a feedback regulatory response.

### The RA robustness response involves autoregulatory changes in the RA metabolic network

To study the RA signaling robustness directly, we developed an experimental protocol to induce transient, physiological disturbances in RA levels. Our study supports the conclusion that the RA metabolic and signaling network exhibits highly efficient robustness to physiological changes. Transcriptome-wide analysis of the restoration of normal gene expression patterns following transient RA manipulation showed that the transcriptomes of RA-manipulated (increased or decreased) and control embryos are highly similar (PCA analysis) at each time point, supporting the robustness of RA signaling to maintain normal, non-teratogenic, target gene expression levels during early gastrulation. A deeper analysis revealed only weak, but measurable, transcriptomic changes that could be attributed to the RA manipulation. Very mild morphological malformations could be observed even when significant changes in RA target genes were observed.

Then, what is the mechanism activated to achieve RA robustness within the physiological range as observed in our study? qPCR analysis provided insights on the RA robustness mechanism showing that target genes exhibited abnormal expression levels at the end of the manipulation (t=0), and some target genes (*hox*) exhibited a speedy return to normal expression after only 1.5 hours from the end of the treatment. Genes encoding RA metabolic network components also exhibited abnormal expression levels at t=0, but their return to normal expression was delayed beyond the time required for the RA targets to reach normal expression. These observations suggest that the RA metabolic network is altered through an RA-dependent feedback regulatory mechanism to restore normal RA signaling and target gene expression.

Multiple reports have described the RA-dependent regulation of individual RA metabolic components. Most studies show up-regulation of enzymes suppressing or reducing RA signaling like CYP26A1, ADHFe1, and DHRS3, or down-regulation of RA producers (anabolic enzymes) like ALDH1A2 and RDH10 as a result of RA treatment (Fujii et al., 1997; Sonneveld et al., 1998; Hollemann et al., 1998; Kam et al., 2013; Dobbs-McAuliffe et al., 2004; Chen et al., 2001; Strate et al., 2009; Sandell et al., 2012; Shabtai et al., 2017). However, many of these studies utilized non-physiological amounts of RA mostly in continuous treatments, they were performed in multiple organisms, cell types, and developmental stages, and some even obtained contradictory results (Dobbs-McAuliffe et al., 2004; Chen et al., 2001; Niederreither et al., 1997; Pavez Loriè et al., 2009; Moss et al., 1998; Hollemann et al., 1998; Fujii et al., 1997; Topletz et al., 2015; Kam et al., 2013; Reijntjes et al., 2010; Strate et al., 2009). Our experimental design based on transcriptomic analysis together with our description of the RA network during early/mid gastrula allowed us to define the RA *network-wide* response to signaling changes, in one organism, at a defined developmental stage, in an unbiased manner by studying all the network components expressed during gastrula *Xenopus* embryos.

We observed that some RA network components exhibit an oscillatory behavior close to control expression levels, probably as a result of the fine-tuning of the RA signal. Such fine-tuning of the RA signal levels coupled to the inherent delay in the transcriptional response could transiently result in the inversion of the overall signaling direction, and an oscillatory transcriptional behavior. In a few instances such paradoxical observations of RA signaling outcome following RA manipulation, i.e. overcompensation, have been reported (Rydeen et al., 2015; D’Aniello et al., 2013; Lee et al., 2012; D’Aniello and Waxman, 2015). These observations identify a very dynamic feedback regulatory network continuously fine-tuning itself to overcome perturbations.

The recovery kinetics analysis after conversion of the RNAseq data to discretized patterns revealed that only 10 out of the 27 possible expression patterns were represented in any of the treatment or control samples, and only five patterns were exhibited by a substantial number of genes (n>100). These observations suggest that RA-regulated genes, direct or indirect, can only exhibit a limited number of possible regulatory responses although intensity differences are possible. Surprisingly, a comparative analysis of the discretized patterns using our unbiased COMPACT approach (Kuttippurathu et al., 2016) revealed that most genes (75.40%) responding to RA manipulation and recovery exhibited a response to only either increased or decreased RA levels. Only 48 genes out of 553 (8.67%) exhibited the commonly expected reciprocal responses to increased versus decreased RA levels. Many of this small group of genes encode RA metabolism or signaling components as part of the robustness response to maintain non-teratogenic RA levels.

The temporal delay in the return to normalcy between RA targets and network components and the network-wide response taking place at multiple levels provide an integrative view of the feedback regulation and robustness response. Initially, the response to RA changes will most probably be managed by the actual enzymes and factors already present in the cell. In parallel, the same RA changes will elicit a self-regulatory transcriptional response which for increased RA should involve the upregulation of RA suppressor activities (DHRS3, ADHFe1, and CYP26A1), and the complementary downregulation of RA producing enzymes (ALDH1A2 and RDH10). This transcriptional response will exhibit a slight delay due to the transcription, translation, and post-translational modifications required.

### Uneven response to increased versus decreased RA levels

Our study provides new insights on the network responses to increased and decreased RA levels. One important conclusion is that these responses are not simple opposites, but are rather under different regulatory rules. As determined from the unbiased discretized pattern analysis, a small proportion of genes respond inversely to increased and decreased RA levels. Otherwise, most of the RA-responsive genes respond to one direction of RA change. This observation was further supported by our relative ranking of the robustness of the different embryo clutches based on the trajectory analysis. We conclude that the RA auto-regulatory robustness response is uneven based on: First, the limited number of RA target genes that respond inversely to increased and decreased RA. Second, the response to reduced signaling is activated in response to even a slight RA reduction, while the response to increased RA is only activated above a threshold. Also, the response to reduced RA reaches an upper response limit, while the response to increased RA appears to have a wider range. Notably, the network responses show different thresholds depending on the direction of the RA manipulation suggesting a high sensitivity to RA reduction but a lower sensitivity to slight RA increase. Third, the alignment of the robustness responses, *hox* changes, fitted a diagonal with a negative slope with a significant correlation. This negative slope suggested that while a clutch might very efficiently deal with increased RA, i.e. high robustness, the same clutch struggles to compensate for a reduction in RA, i.e. low robustness, suggesting an intrinsic trade-off along the lines of a Pareto multiobjective optimization (Miettinen, 1998; Tendler et al., 2015; Schuetz et al., 2012; Shoval et al., 2012). In the present case, the Pareto optimality suggests that improving robustness towards one objective (e.g., decreased RA) can only be achieved by degrading robustness towards another (increased RA). Our results showing that some clutches achieve only moderate robustness to positive and negative fluctuations in RA, whereas others show uneven robustness, are consistent with the notion of Pareto optimality of robustness to RA fluctuations in the early embryo. Fourth, different clutches with similar robustness can mount robustness responses by incorporating different components of the RA network. The transcriptomic and HT-qPCR analyses allowed us to perform a detailed determination of the network components comprising a robustness response across different clutches to uncover its mechanism. The RA metabolic network includes multiple enzymes with overlapping biochemical activities, e.g., ALDH1A1 (RALDH1), ALDH1A2 (RALDH2), and ALDH1A3 (RALDH3) enzymes that oxidize retinaldehyde to produce RA (Kedishvili, 2016, 2013; Shabtai et al., 2016; Ghyselinck and Duester, 2019). The enzymatic efficiencies and expression patterns including location, timing, and intensity may be different across the enzymes (Blentic et al., 2003; Lupo et al., 2005; Chen et al., 2001; Romand et al., 2004; Shabtai et al., 2016). The enzymes CYP26A1, B1, and C1, or DHRS3, ADHFe1, and RDH14 function to prevent excessive RA signaling (Belyaeva et al., 2008, 2017; Shabtai et al., 2017; Sonneveld et al., 1998; Hollemann et al., 1998). These redundant enzymes and factors can help establish a robustness response using different components (Figs. 8D,E and 9). Finally, genetic polymorphisms can probably partially explain the different responses between clutches of our outbred experimental model, *Xenopus laevis* (Savova et al., 2017). Formally, technical issues could account for some variability. The six clutches analyzed by HT-qPCR, six clutches analyzed by RNAseq, and the five clutches from the qPCR analysis showed a similar distribution of the same variability supporting the possible involvement of genetic polymorphisms. We mined the Savova et al. (2017) data identifying multiple polymorphisms in RA network components (Supplemental Table S3), supporting the potential contribution of genetic variability to the differential and uneven robustness across clutches.

### Retinoic acid signaling changes due to environmental changes and disease risk

While RA exposure induces dramatic developmental malformations, treatment of the same embryos with ROL or RAL, the RA precursors, requires higher concentrations (70-100X and 5X respectively) to induce similar defects to the RA treatment (Durston et al., 1989; Yelin et al., 2005). The important biochemical difference between the compounds is that ROL or RAL have to go through RA biosynthesis to become the active ligand, while RA is already the final product. The oxidation of ROL to RAL is a reversible reaction that can reduce substrate availability for the RALDH enzymes, while excess RA can only be countered through inactivation by retinoic acid hydroxylating enzymes, e.g., CYP26 (Sakai et al., 2001; Hollemann et al., 1998; Dobbs-McAuliffe et al., 2004). Therefore, the reduced teratogenicity of the precursors could be the result of reduced conversion to RA as part of the feedback regulation of this network, i.e. robustness, while the robustness response is more restricted in its regulatory feedback management of RA addition.

All our experiments were performed by either partially inhibiting the endogenous levels of RA, or by slightly increasing (∼10% increase) the physiological RA content. Under these conditions, the RA robustness of the embryo efficiently normalizes the transcriptome via regulatory feedback. Based on the comparative analysis of 12 clutches (genetic backgrounds), we suggest that robustness efficiency will have a threshold beyond which it will become ineffective in restoring normal RA signaling. Our clutch analysis suggests that this threshold might be strongly dependent on genetic polymorphisms affecting enzymatic activity or gene expression parameters. Consistent with these observations, a threshold or toxicological tipping point for RA signaling was recently described in a cell-based model (Saili et al., 2019). The clutch analysis also suggested that the RA network response might also be dependent on genetic variability like promoter polymorphisms. Then, the developmental malformations arising from environmental insults on RA signaling largely depend on genetic polymorphisms which will determine the composition, efficiency and threshold of the response and the actual network components comprising such a response.

### AVAILABILITY OF SUPPORTING DATA/ADDITIONAL FILES

The raw and normalized datasets for the RNAseq and HT-qPCR data are available online as Gene Expression Omnibus datasets via SuperSeries GSE154408 containing RNAseq data: GSE154399 and HT-qPCR data: GSE154407.

## Supporting information

Supplemental Data 1 COMPACT

Supplemental Data 2 WGCNA

## ABBREVIATIONS USED

COMPACT: Comparative Matrix of Pattern Counts

DEAB: 4-diethylaminobenzaldehyde

PCA: Principal Component Analysis

PCR: Polymerase Chain Reaction

RA: retinoic acid

WGCNA: Weighted Gene Coexpression Network Analysis

## AUTHOR CONTRIBUTIONS

R.V. and A.F. conceived and supervised the study and designed the experiments and analysis methodology. L.B-K., M.G., T.A., K.K., and A.F. performed embryo experiments and real-time PCR assessment and developed the figures. A.B. performed the initial analysis of the RNAseq data. S.A. conducted the high-throughput PCR validation of RNAseq results. M.P. conducted the analysis of transcriptomics and HT-qPCR data and performed the network and trajectory analyses and developed the figures. M.P., R.V., and A.F. interpreted the results and drafted the manuscript.

---

**Supplemental Figure Sl.**
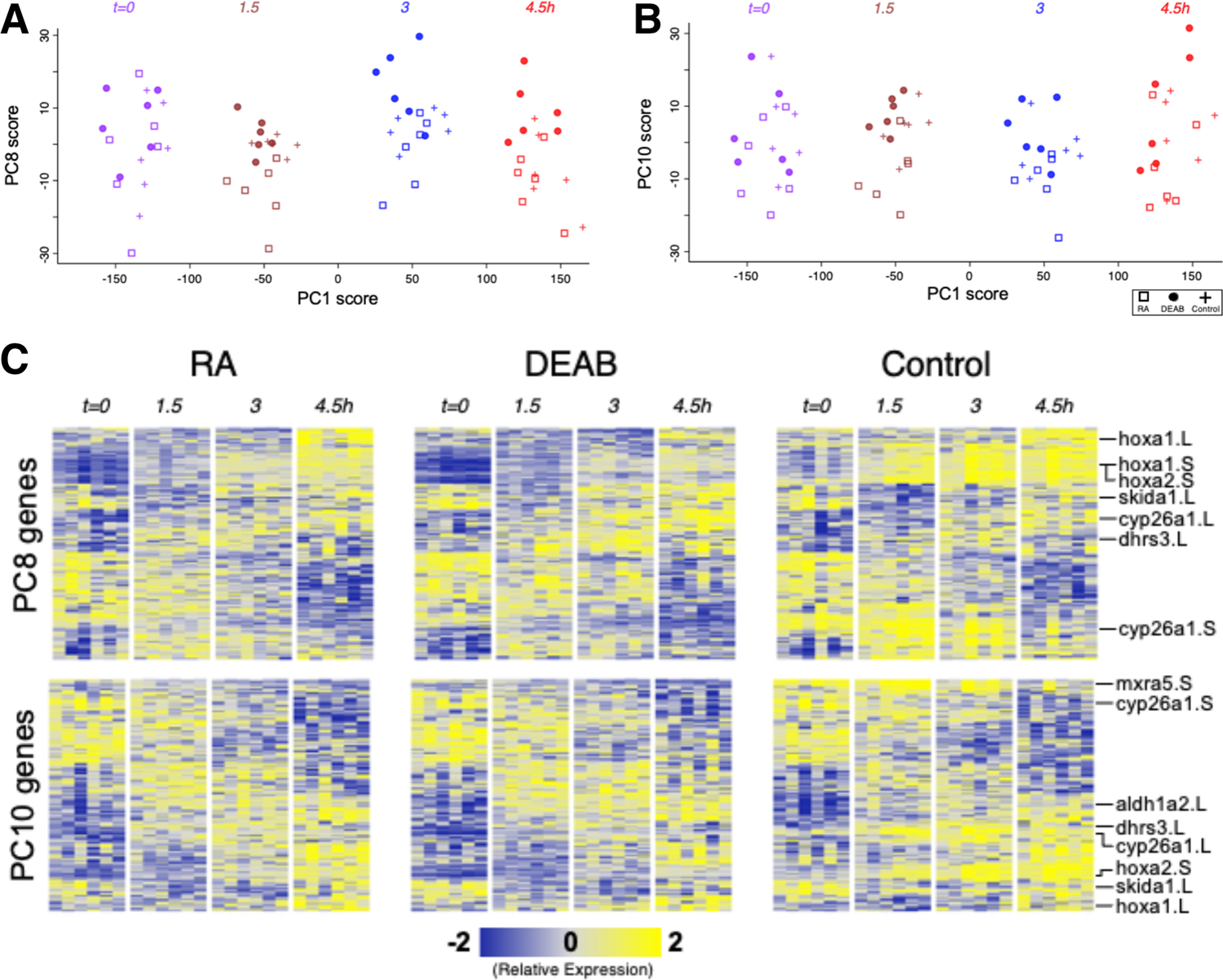
Efficient RA signaling robustness at the transcriptomic level. **(A,B)** Principal Component Analysis of six clutches over time for **(A)** PC1/PC8, and **(B)** PCl/PClO. **(C)** Heatmap of gene expression of the top-100 positive and top-100 negative loadings corresponding to PCS and PClO. A subset of the genes is highlighted based on the relevance to early developmental process.

**Supplemental Figure S2.**
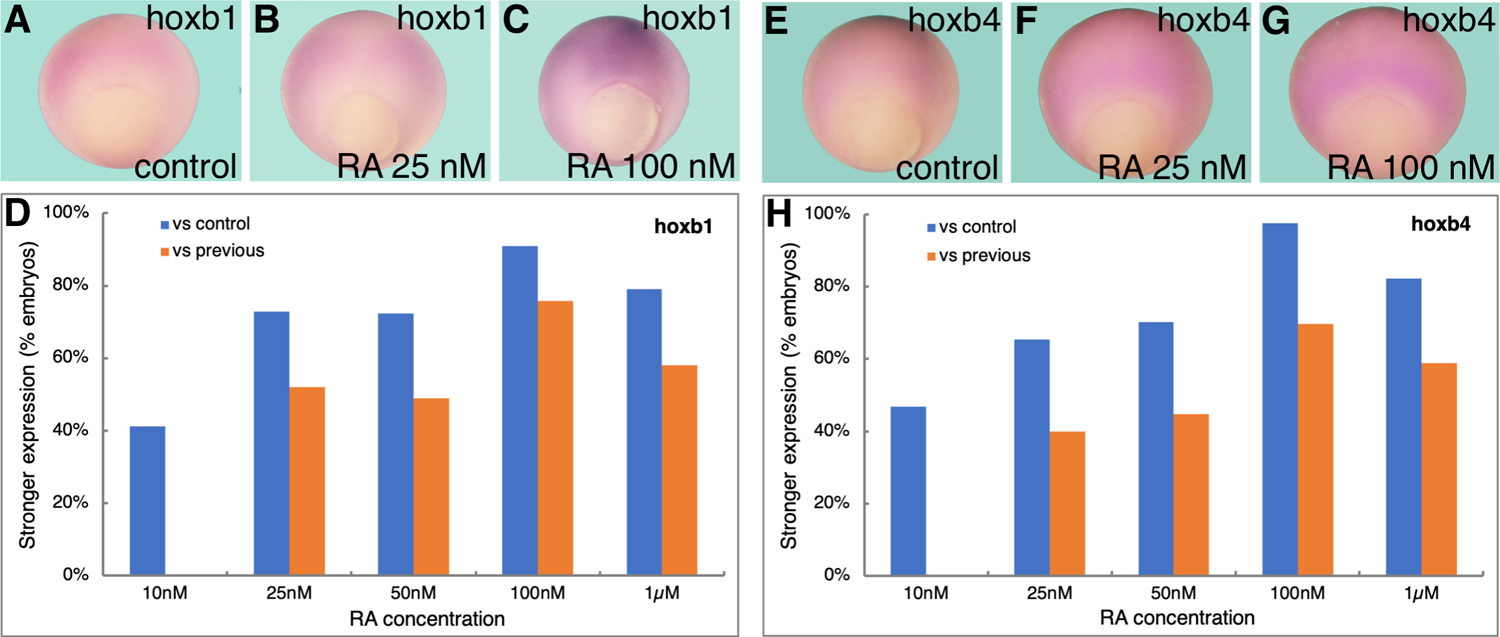
Dose response of *hoxbl* and *hoxb4* to increased retinoic acid levels. Embryos were treated with increased retinoic acid concentrations (10 nM - 1 µM) until mid gastrula (st. 11) and then processed for *in situ* hybridization with probes specific for *hoxbl* (A-C) or *hoxb4* **(E-G). (D,H)** The changes in the expression pattern for both genes was scored as an increase in signal intensity from the control group (vs. control) or the previous lower concentration (vs. previous).

**Supplemental Figure S3.**
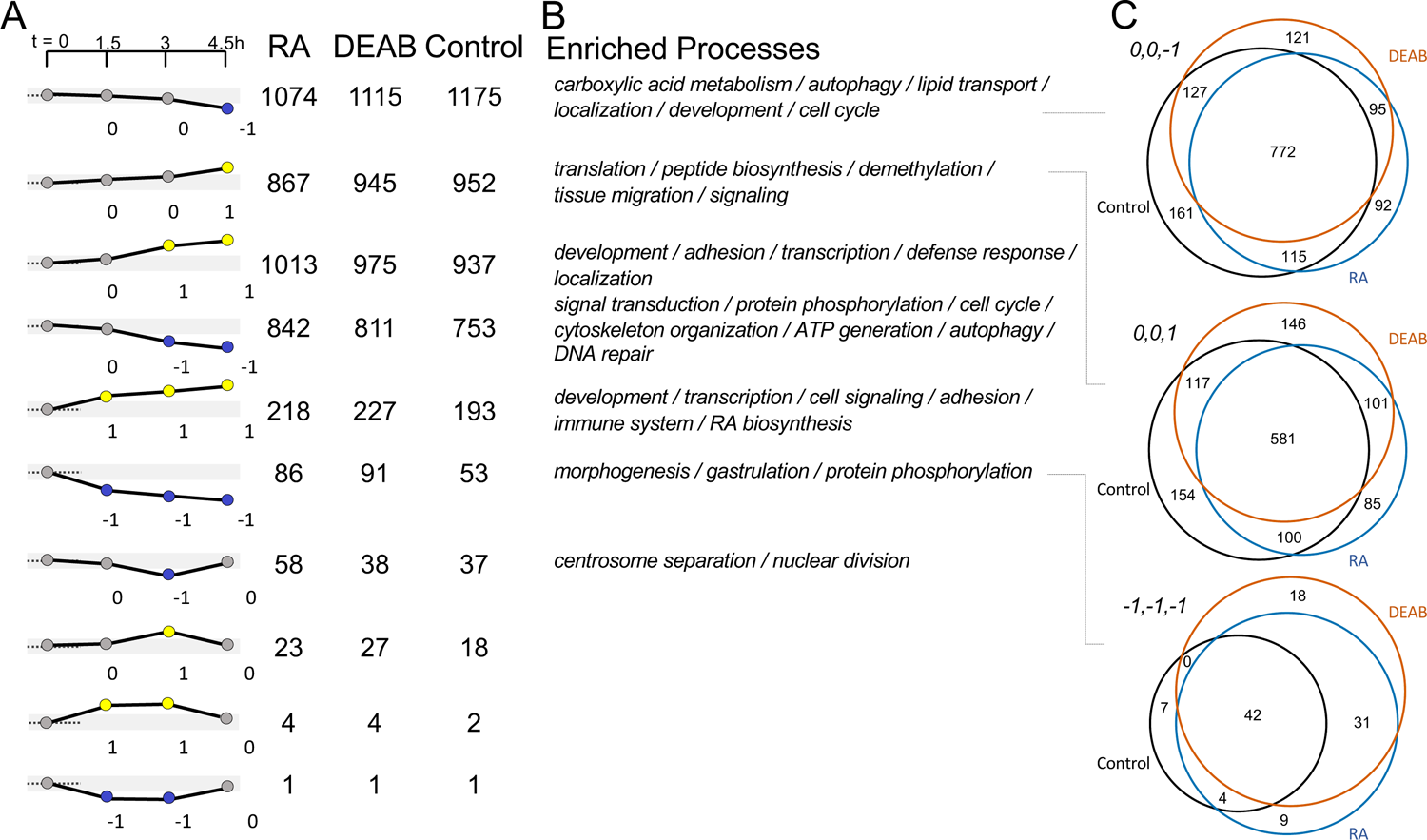
Transcriptome overlap between RA manipulation and controls. **(A)** Genes were grouped into the 27 possible discretized expression patterns based on up-regulation (yellow), down-regulation (blue), and no change (grey) above the 2-fold change threshold at each of the three recovery time points compared to the t=0 sample. The ten dynamic patterns that contained at least one gene are shown. The numbers below each graph exemplify the discretized pattern as a numeric vector (+1, 0, −1). The counts next to the patterns indicate the number of genes that show the corresponding expression pattern in each of the two treatment groups and the controls. **(B)** The Gene Ontology biological processes statistically enriched in the control group are indicated alongside the pattern counts. Details of statistical analysis results are available in Supplemental Table S1. **(C)** Venn diagrams to compare the overlap between the two treatments and control, illustrated for three differential gene expression patterns. The number of genes that showed similar differential expression patterns in one or more experimental groups are indicated in the corresponding overlapping regions. black - control; orange - DEAB; blue - RA.

**Supplemental Figure S4.**
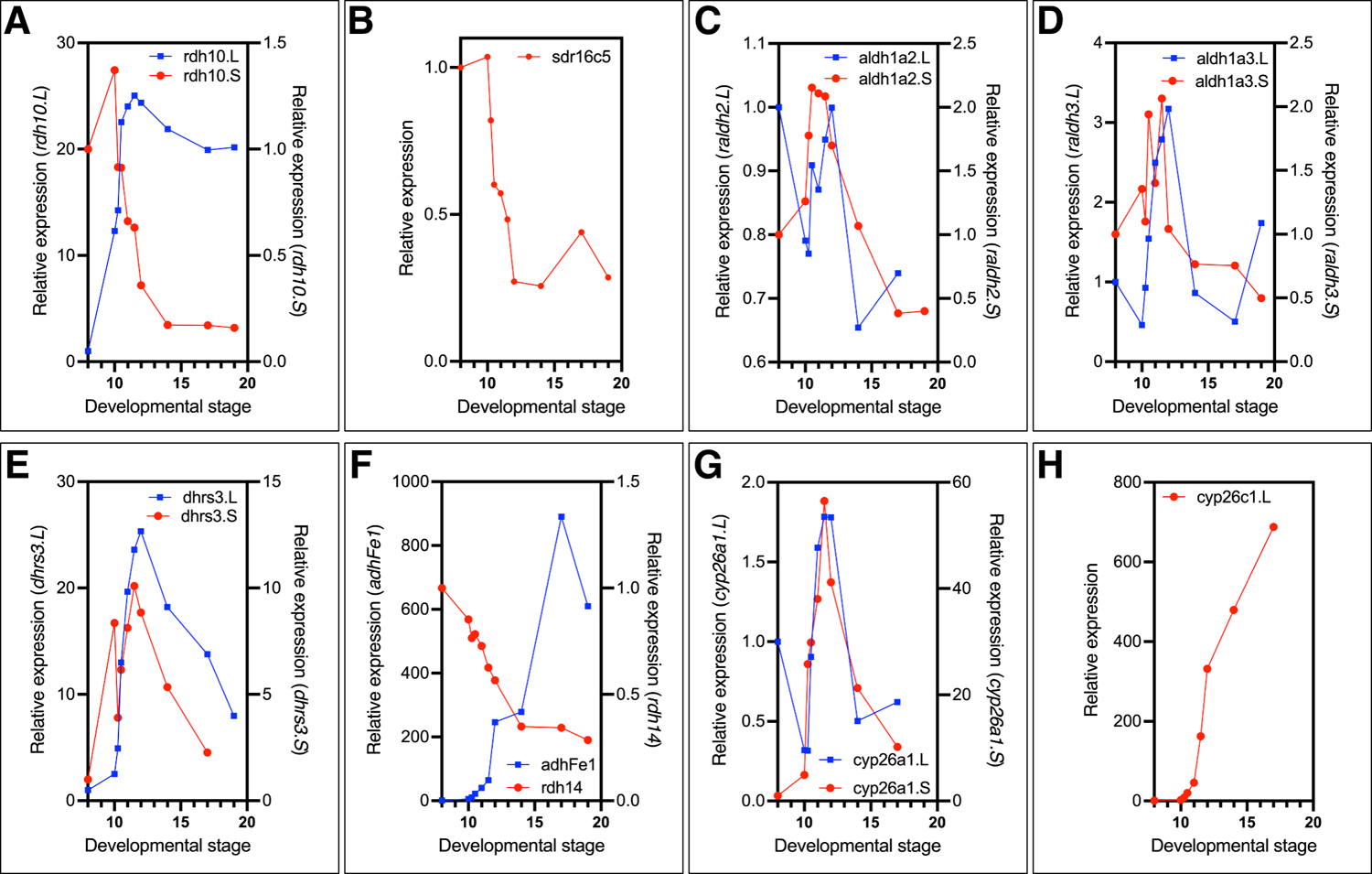
Temporal expression pattern of the main RA network components studied. To determine the temporal pattern of expression of the RA network components, RNA samples from embryos from midblastula (st. 8) to advanced neurula stages (st. 19) were collected. Relative expression was determined by qPCR for: **(A)** *rdhlO.L, rdhlO.S;* **(B)** *sdr16c5;* **(C)** *aldhla2.L, aldhla2.S;* **(D)** *aldhla3.L, aldhla3.S;* **(E)** *dhrs3.L, dhrs3.S;* **(F)** *adhFel, rdh14;* **(G)** *cyp26al .L, cyp26al.S;* **(H)** *cyp26cl .L*.

**Supplemental Figure S5.**
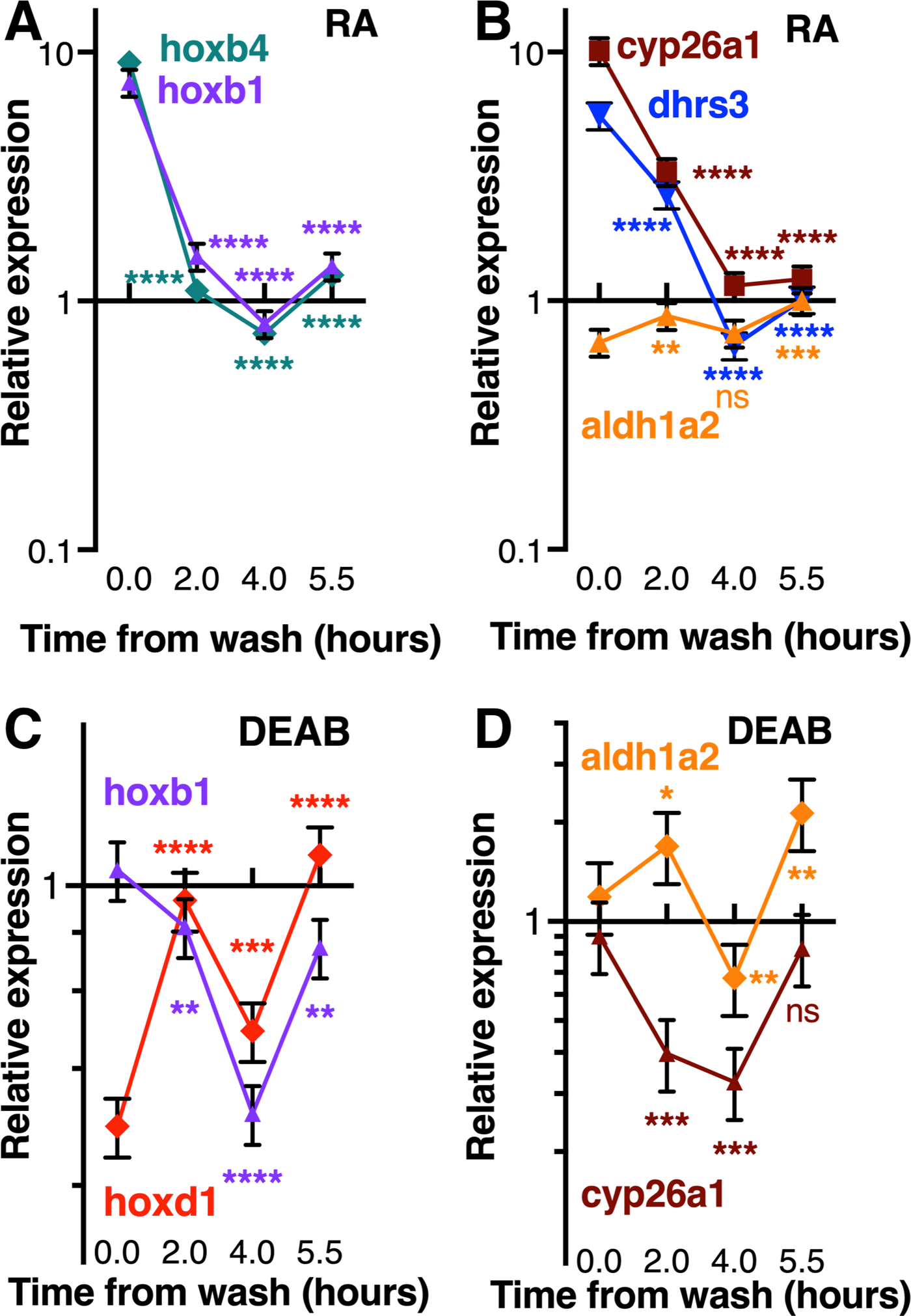
Kinetics of the recovery from RA manipulation. Embryos were transiently treated with either 10 nM RA **(A,B),** or 50 µM DEAB **(C,D).** Treatments were initiated during late blastula (st. 9.5), and washed by early gastrula (st. 10.25). RNA samples were collected at different time points during the recovery period. The response of genes, RA target, and RA metabolic enzymes was studied by qPCR. Statistical significance (Student’s t-test) was calculated compared to the expression at the end of the treatment (tO). *, p<0.05; **, p<0.01; ***, p<0.001; ****, p<0.0001; ns, not significant.

**Supplemental Figure S6.**
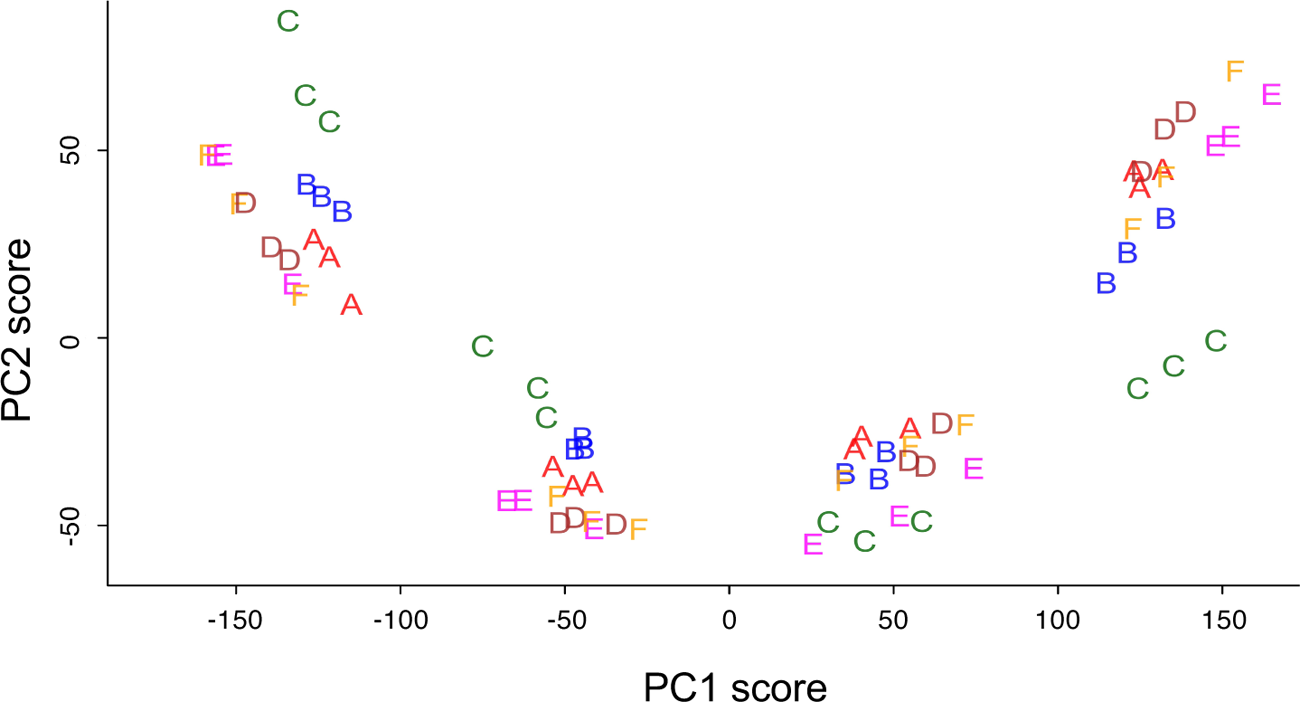
Principal Component Analysis suggests clutch-to-clutch variation. Labeling of the different clutches reveals clustering of all samples of the same clutch at all time points analyzed.

**Supplemental Figure S7.**
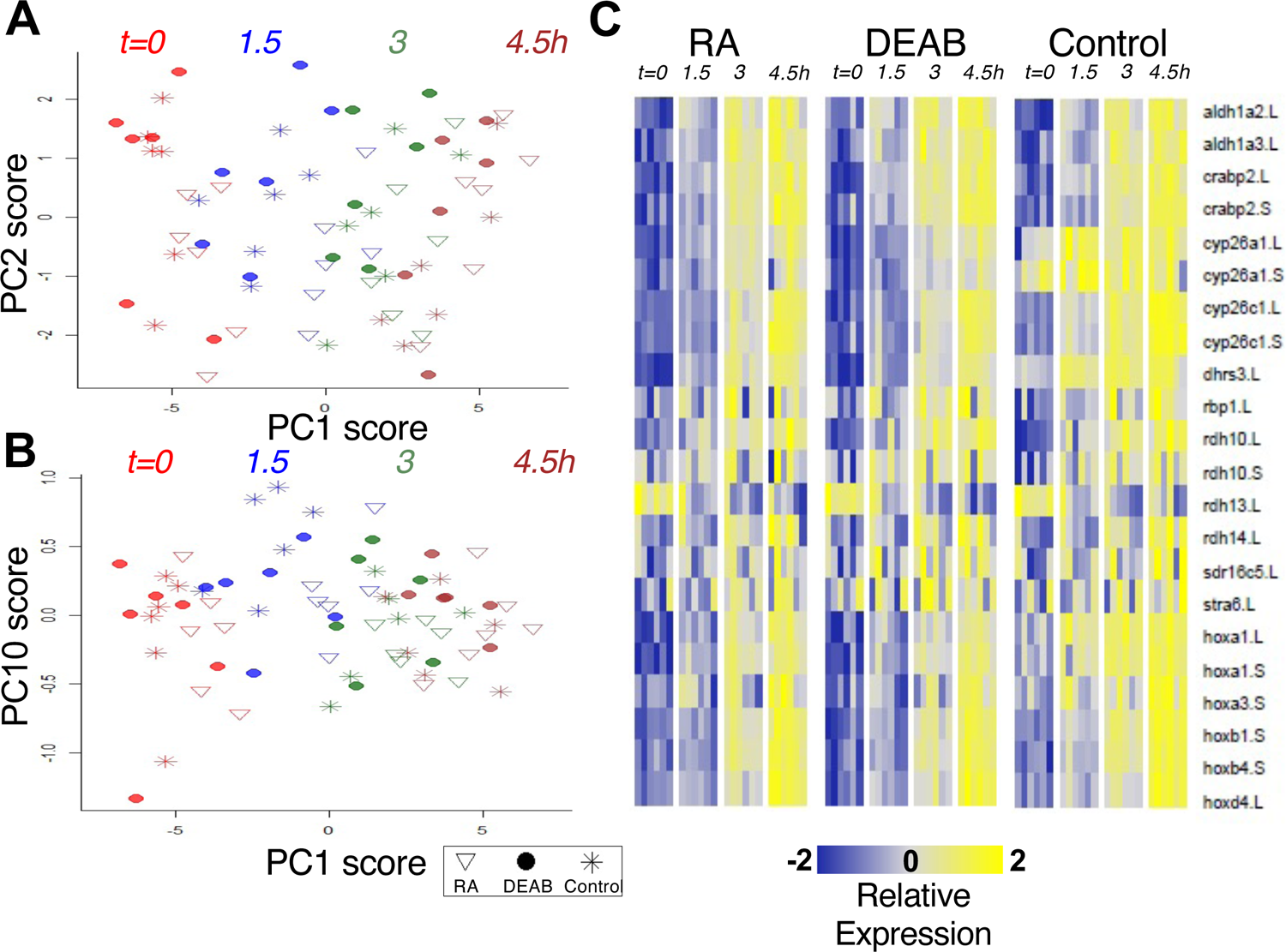
HT-qPCR analysis of RA network components. Principal Component Analysis of the HT-qPCR data revealed heterogeneity across developmental stages and treatments. **(A)** Distribution of samples along PC1 and PC2 axes. **(B)** Distribution of samples along PC1 and PC10 axes. **(C)** Heatmap of time series differential expression of RA network genes.

**Supplemental Figure S8.**
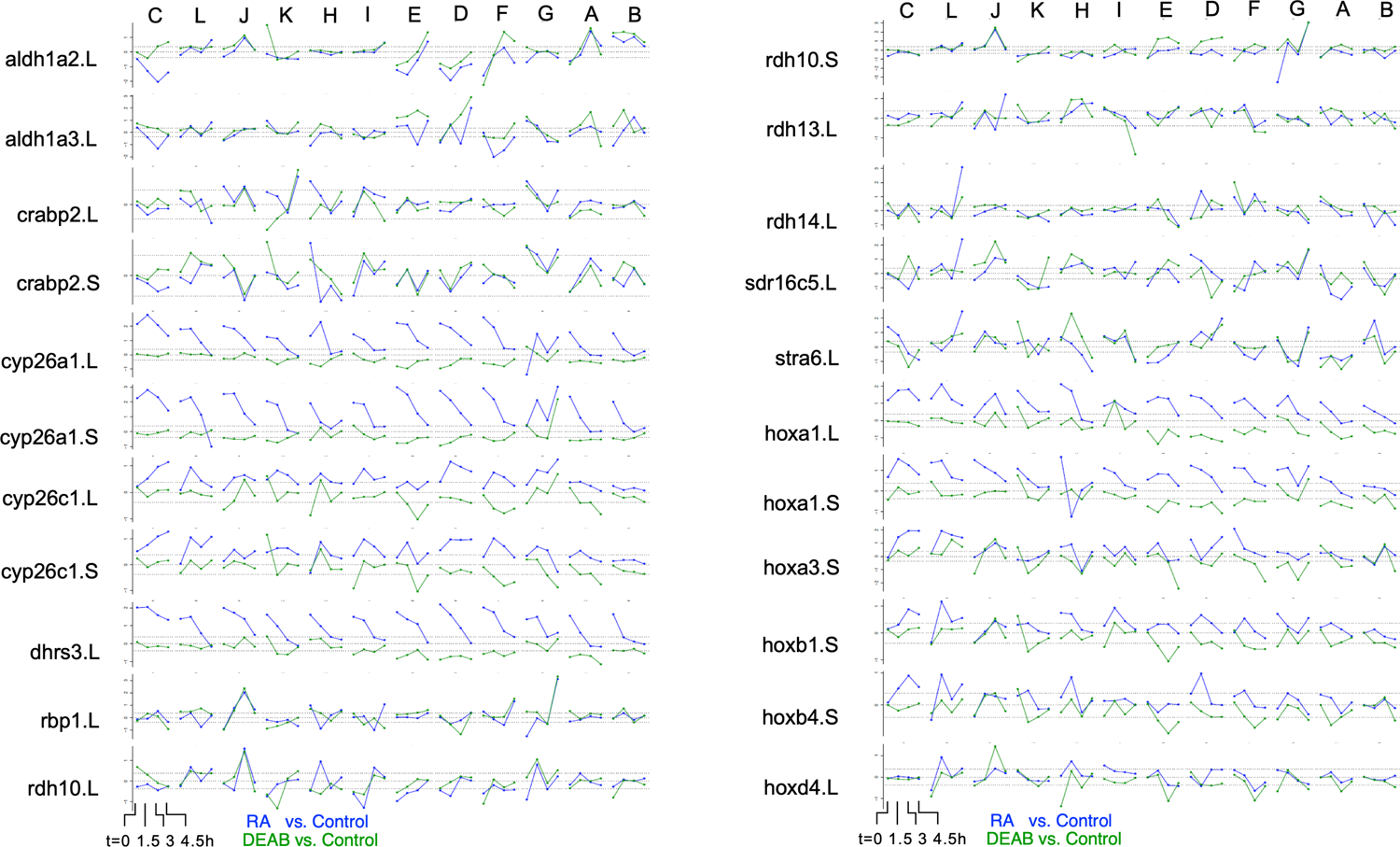
Clutch-wise heterogeneity in the response of RA network genes following RA manipulation. Clutch-wise differential expression dynamics of RA network genes based on the RNAseq and HT-qPCR data. The clutches are ordered left to right according to the time taken for recovery of *hoxal.L* expression.

**Supplemental Figure S9.**
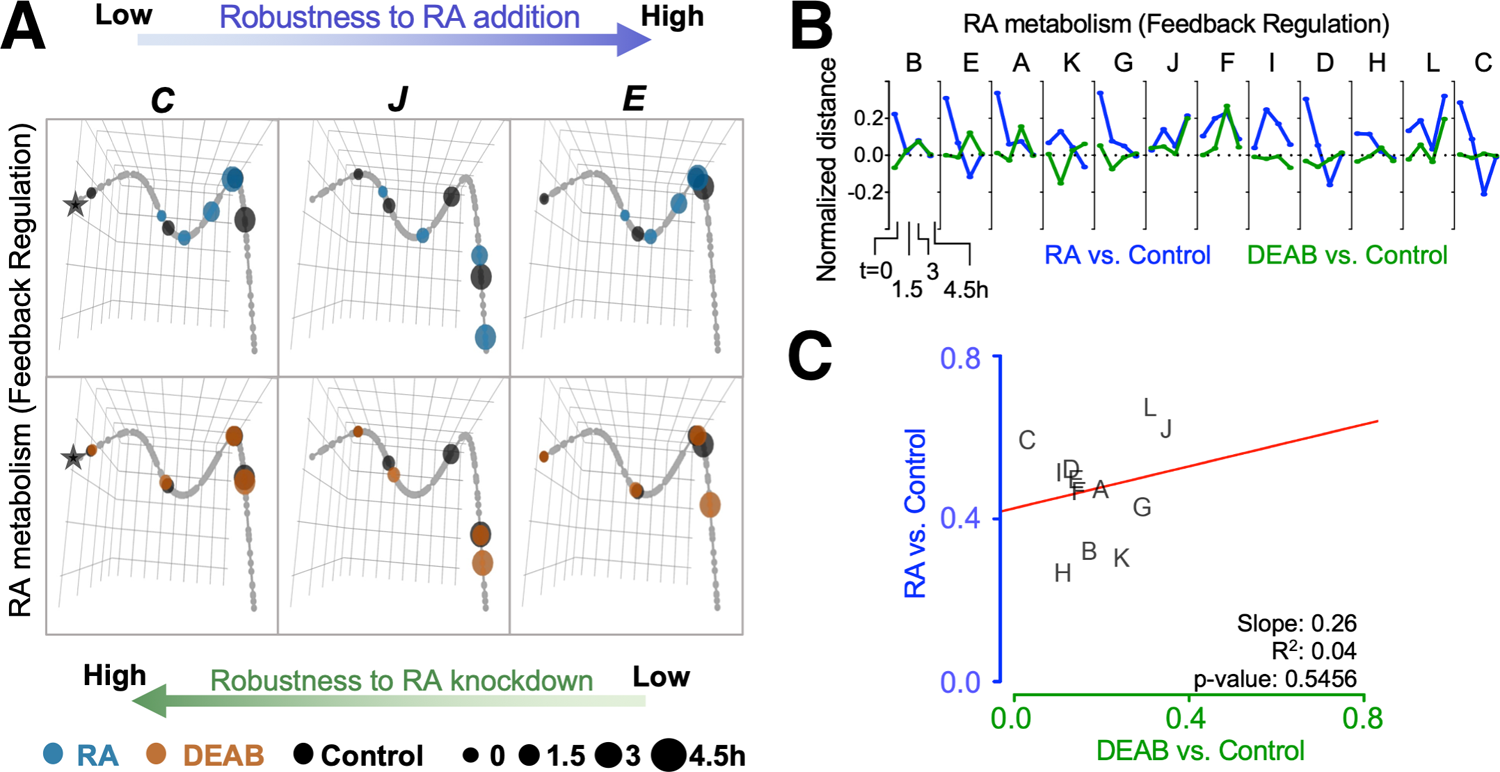
Trajectory analysis to compare the extent of individual clutch robustness based on RA network component expression. **(A)** 3-dimensional principal curve for the RA network genes, showing projections of the sample points on the curve for **(A, top)** RA and **(A, bottom)** DEAB treatments. Principal curves for clutches C, J, and E are shown. The black star indicates the beginning of the curve for the distance measurement along the trajectory. Ranking of clutches is based on the net (absolute) normalized distance of treatment samples from the corresponding Control sample for each time point. **(B)** Normalized expression shift profile calculated from the principal curve as the arc distance between the treatment and the corresponding control. Clutches are rank­ ordered from lowest to highest net expression shift for *hox* genes in the RA group. Clutches A-F data from RNA­ seq, clutches G-L data from HT-qPCR. (C) Distribution of clutches based on the trajectory determined robustness to RA and DEAB treatments relative to each other. The letters indicate the distinct clutches.

**Supplemental Table S1.** List of Gene Ontology annotations with corresponding genes and statistical significance corresponding to Supplemental Figure S3.

| Control Pattern | ID | Description | Total Genes | Pathway Genes | p value | adjusted q value |
| --- | --- | --- | --- | --- | --- | --- |
| 0 0 0 -1 | GO:0046626 | regulation of insulin receptor signaling pathway | 426 | 5 | 2.90E-06 | 3.99E-03 |
|  | GO:0006914 | autophagy | 426 | 14 | 7.65E-05 | 1.50E-02 |
|  | GO:0006631 | fatty acid metabolic process | 426 | 11 | 1.45E-04 | 2.24E-02 |
|  | GO:0006096 | glycolytic process | 426 | 7 | 5.28E-04 | 3.80E-02 |
|  | GO:0008354 | germ cell migration | 426 | 3 | 7.55E-04 | 4.02E-02 |
|  | GO:0009799 | specification of symmetry | 426 | 5 | 1.04E-03 | 4.74E-02 |
|  | GO:0010876 | lipid localization | 426 | 13 | 1.07E-03 | 4.74E-02 |
| 0 0 0 1 | GO:0006518 | peptide metabolic process | 302 | 44 | 5.82E-13 | 2.24E-10 |
|  | GO:0001708 | cell fate specification | 302 | 4 | 2.34E-03 | 2.11E-01 |
|  | GO:0060485 | mesenchyme development | 302 | 7 | 3.95E-03 | 2.68E-01 |
|  | GO:0090130 | tissue migration | 302 | 3 | 8.10E-03 | 4.42E-01 |
|  | GO:0010631 | epithelial cell migration | 302 | 2 | 8.83E-03 | 4.42E-01 |
| 0 0 1 1 | GO:0048513 | animal organ development | 320 | 35 | 2.34E-08 | 1.95E-05 |
|  | GO:0007155 | cell adhesion | 320 | 26 | 3.21E-07 | 7.75E-05 |
|  | GO:0030334 | regulation of cell migration | 320 | 5 | 6.53E-03 | 7.35E-02 |
|  | GO:1905209 | positive regulation of cardiocyte differentiation | 320 | 2 | 9.88E-03 | 8.84E-02 |
| 0 0 -1 -1 | GO:0007020 | microtubule nucleation | 250 | 3 | 3.01E-03 | 4.81E-01 |
|  | GO:0009967 | positive regulation of signal transduction | 250 | 9 | 5.54E-03 | 4.81E-01 |
|  | GO:0098927 | vesicle-mediated transport between endosomal compartments | 250 | 2 | 6.11E-03 | 4.81E-01 |
|  | GO:0010256 | endomembrane system organization | 250 | 7 | 6.13E-03 | 4.81E-01 |
| 0 1 1 1 | GO:0048513 | animal organ development | 83 | 15 | 5.49E-07 | 1.25E-04 |
|  | GO:0048732 | gland development | 83 | 5 | 1.42E-06 | 1.25E-04 |
|  | GO:0010557 | positive regulation of macromolecule biosynthetic process | 83 | 11 | 3.29E-06 | 1.97E-04 |
|  | GO:0031328 | positive regulation of cellular biosynthetic process | 83 | 11 | 4.39E-06 | 1.97E-04 |
|  | GO:0016055 | Wnt signaling pathway | 83 | 8 | 2.71E-04 | 4.57E-03 |
|  | GO:0044344 | cellular response to fibroblast growth factor stimulus | 83 | 4 | 4.94E-04 | 6.21E-03 |
|  | GO:0010817 | regulation of hormone levels | 83 | 4 | 1.92E-03 | 1.84E-02 |
|  | GO:0042572 | retinol metabolic process | 83 | 2 | 2.43E-03 | 2.28E-02 |
| 0 -1 -1 -1 | GO:0008285 | negative regulation of cell population proliferation | 15 | 3 | 7.57E-05 | 6.61E-03 |
|  | GO:0007369 | gastrulation | 15 | 2 | 5.21E-03 | 9.93E-02 |
| 0 0 -1 0 | GO:0007100 | mitotic centrosome separation | 14 | 2 | 1.87E-05 | 2.16E-03 |
|  | GO:0140014 | mitotic nuclear division | 14 | 2 | 5.71E-03 | 8.00E-02 |
|  | GO:0051128 | regulation of cellular component organization | 14 | 3 | 8.47E-03 | 8.00E-02 |
|  | GO:0031338 | regulation of vesicle fusion | 14 | 1 | 8.51E-03 | 8.00E-02 |
|  | GO:0032418 | lysosome localization | 14 | 1 | 9.92E-03 | 8.00E-02 |
| 0 0 1 0 | GO:0045663 | positive regulation of myoblast differentiation | 4 | 1 | 2.84E-03 | 1.71E-02 |
|  | GO:0045445 | myoblast differentiation | 4 | 1 | 3.25E-03 | 1.71E-02 |

**Supplemental Table S2.**
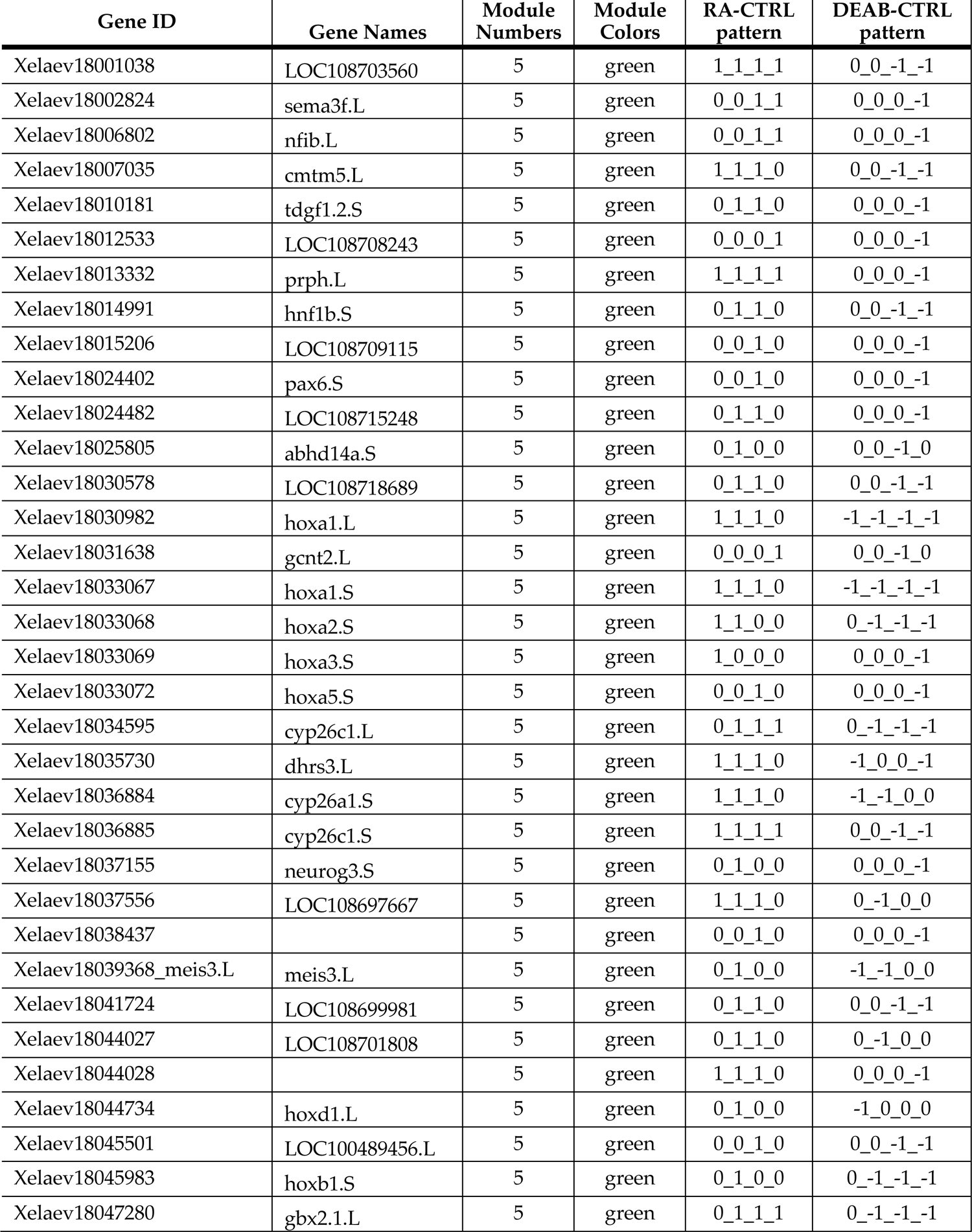
Weighted Gene Correlation Network Analysis (WGCNA)

**Supplemental Table S3.**
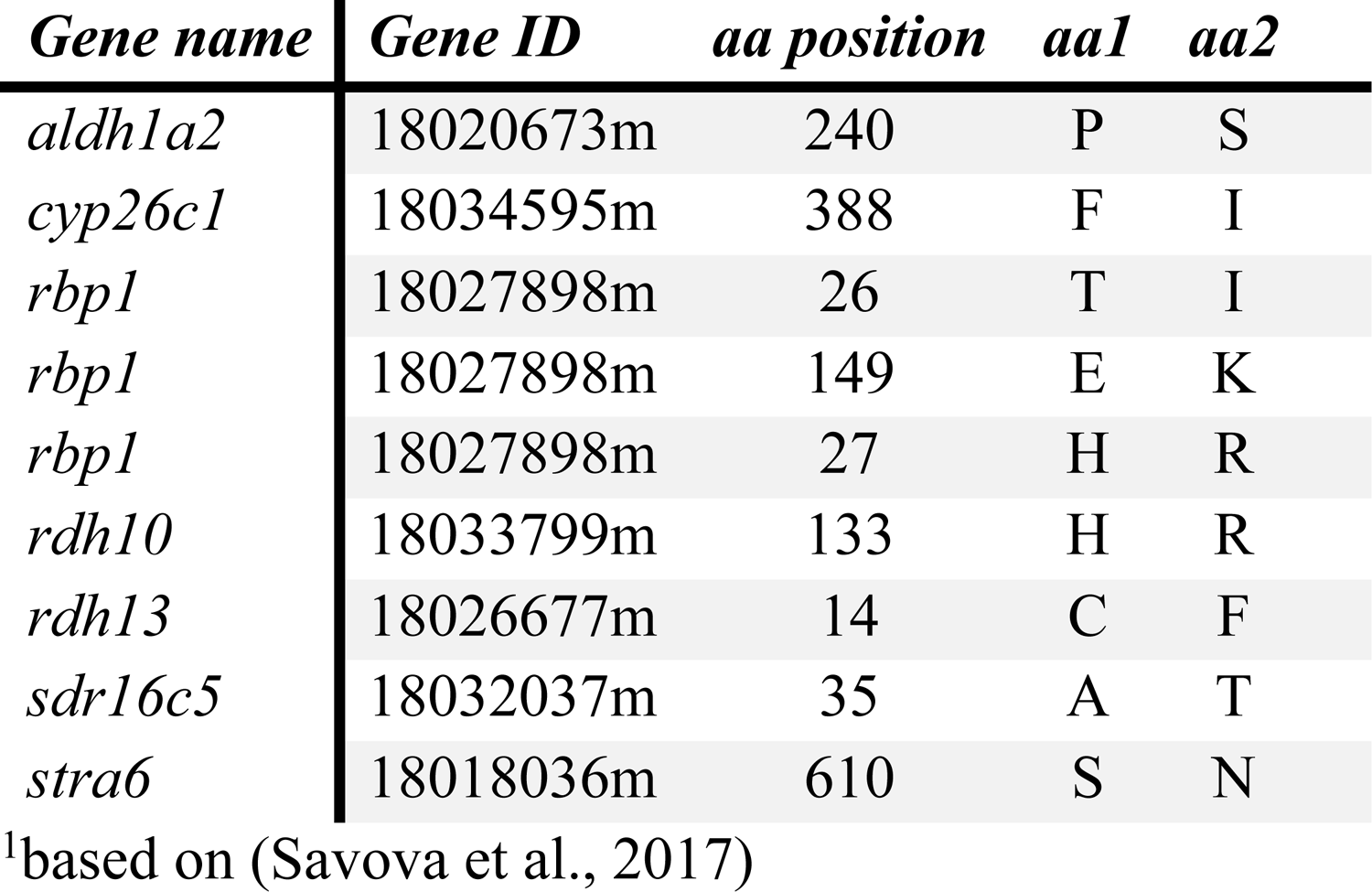
RA network component polymorphisms between *Xenopus* strains^1^

## BIBLIOGRAPHY

1. Adams, M. K., Belyaeva, O. V., Wu, L., and Kedishvili, N. Y. (2014). The retinaldehyde reductase activity of DHRS3 is reciprocally activated by retinol dehydrogenase 10 to control retinoid homeostasis. J. Biol. Chem. 289, 14868– 14880. doi:10.1074/jbc.M114.552257.

2. Begemann, G., Schilling, T. F., Rauch, G. J., Geisler, R., and Ingham, P. W. (2001). The zebrafish neckless mutation reveals a requirement for raldh2 in mesodermal signals that pattern the hindbrain. Development 128, 3081–3094.

3. Belyaeva, O. V., Adams, M. K., Wu, L., and Kedishvili, N. Y. (2017). The antagonistically bifunctional retinoid oxidoreductase complex is required for maintenance of all-trans-retinoic acid homeostasis. J. Biol. Chem. 292, 5884– 5897. doi:10.1074/jbc.M117.776914.

4. Belyaeva, O. V., Korkina, O. V., Stetsenko, A. V., and Kedishvili, N. Y. (2008). Human retinol dehydrogenase 13 (RDH13) is a mitochondrial short-chain dehydrogenase/reductase with a retinaldehyde reductase activity. FEBS J. 275, 138–147. doi:10.1111/j.1742-4658.2007.06184.x.

5. Billings, S. E., Pierzchalski, K., Butler Tjaden, N. E., Pang, X.-Y., Trainor, P. A., Kane, M. A., and Moise, A. R. (2013). The retinaldehyde reductase DHRS3 is essential for preventing the formation of excess retinoic acid during embryonic development. FASEB J. 27, 4877–4889. doi:10.1096/fj.13-227967.

6. Blaner, W. S., Li, Y., Brun, P.-J., Yuen, J. J., Lee, S.-A., and Clugston, R. D. (2016). Vitamin A absorption, storage and mobilization. Subcell Biochem 81, 95–125. doi:10.1007/978-94-024-0945-1_4.

7. Blaner, W. S. (2019). Vitamin A signaling and homeostasis in obesity, diabetes, and metabolic disorders. Pharmacol. Ther. 197, 153–178. doi:10.1016/j.pharmthera.2019.01.006.

8. Blentic, A., Gale, E., and Maden, M. (2003). Retinoic acid signalling centres in the avian embryo identified by sites of expression of synthesising and catabolising enzymes. Dev. Dyn. 227, 114–127. doi:10.1002/dvdy.10292.

9. Blumberg, B., Bolado, J., Moreno, T. A., Kintner, C., Evans, R. M., and Papalopulu, N. (1997). An essential role for retinoid signaling in anteroposterior neural patterning. Development 124, 373–379.

10. Carlson, M. (2017). Genome wide annotation for Xenopus. Bioconductor. doi:10.18129/b9.bioc.org.xl.eg.db.

11. Chen, Y., Huang, L., and Solursh, M. (1994). A concentration gradient of retinoids in the early Xenopus laevis embryo. Dev. Biol. 161, 70–76. doi:10.1006/dbio.1994.1008.

12. Chen, Y., Pollet, N., Niehrs, C., and Pieler, T. (2001). Increased XRALDH2 activity has a posteriorizing effect on the central nervous system of Xenopus embryos. Mech. Dev. 101, 91–103. doi:10.1016/S0925-4773(00)00558-X.

13. Corcoran, J., So, P. L., and Maden, M. (2002). Absence of retinoids can induce motoneuron disease in the adult rat and a retinoid defect is present in motoneuron disease patients. J. Cell Sci. 115, 4735–4741.

14. Creech Kraft, J., Kimelman, D., and Juchau, M. R. (1995). Xenopus laevis: a model system for the study of embryonic retinoid metabolism. II. Embryonic metabolism of all-trans-3,4-didehydroretinol to all-trans-3,4-didehydroretinoic acid. Drug Metab. Dispos. 23, 83–89.

15. Creech Kraft, J., Schuh, T., Juchau, M. R., and Kimelman, D. (1994). Temporal distribution, localization and metabolism of all-trans-retinol, didehydroretinol and all-trans-retinal during Xenopus development. Biochem. J. 301 ( Pt 1), 111–119.

16. Cui, J., Michaille, J.-J., Jiang, W., and Zile, M. H. (2003). Retinoid receptors and vitamin A deficiency: differential patterns of transcription during early avian development and the rapid induction of RARs by retinoic acid. Dev. Biol. 260, 496–511.

17. D’Aniello, E., Rydeen, A. B., Anderson, J. L., Mandal, A., and Waxman, J. S. (2013). Depletion of retinoic acid receptors initiates a novel positive feedback mechanism that promotes teratogenic increases in retinoic acid. PLoS Genet. 9, e1003689. doi:10.1371/journal.pgen.1003689.

18. D’Aniello, E., and Waxman, J. S. (2015). Input overload: Contributions of retinoic acid signaling feedback mechanisms to heart development and teratogenesis. Dev. Dyn. 244, 513–523. doi:10.1002/dvdy.24232.

19. Dobbs-McAuliffe, B., Zhao, Q., and Linney, E. (2004). Feedback mechanisms regulate retinoic acid production and degradation in the zebrafish embryo. Mech. Dev. 121, 339–350. doi:10.1016/j.mod.2004.02.008.

20. Durston, A. J., Timmermans, J. P., Hage, W. J., Hendriks, H. F., de Vries, N. J., Heideveld, M., and Nieuwkoop, P. D. (1989). Retinoic acid causes an anteroposterior transformation in the developing central nervous system. Nature 340, 140–144. doi:10.1038/340140a0.

21. Eldar, A., Shilo, B.-Z., and Barkai, N. (2004). Elucidating mechanisms underlying robustness of morphogen gradients. Curr. Opin. Genet. Dev. 14, 435–439. doi:10.1016/j.gde.2004.06.009.

22. Epstein, M., Pillemer, G., Yelin, R., Yisraeli, J. K., and Fainsod, A. (1997). Patterning of the embryo along the anterior-posterior axis: the role of the caudal genes. Development 124, 3805–3814.

23. Fainsod, A., Bendelac-Kapon, L., and Shabtai, Y. (2020). Fetal alcohol spectrum disorder: embryogenesis under reduced retinoic acid signaling conditions. Subcell Biochem 95, 197–225. doi:10.1007/978-3-030-42282-0_8.

24. Fainsod, A., and Kot-Leibovich, H. (2018). Xenopus embryos to study fetal alcohol syndrome, a model for environmental teratogenesis. Biochem. Cell Biol. 96, 77– 87. doi:10.1139/bcb-2017-0219.

25. Feng, L., Hernandez, R. E., Waxman, J. S., Yelon, D., and Moens, C. B. (2010). Dhrs3a regulates retinoic acid biosynthesis through a feedback inhibition mechanism. Dev. Biol. 338, 1–14. doi:10.1016/j.ydbio.2009.10.029.

26. Fujii, H., Sato, T., Kaneko, S., Gotoh, O., Fujii-Kuriyama, Y., Osawa, K., Kato, S., and Hamada, H. (1997). Metabolic inactivation of retinoic acid by a novel P450 differentially expressed in developing mouse embryos. EMBO J. 16, 4163–4173. doi:10.1093/emboj/16.14.4163.

27. Ghyselinck, N. B., and Duester, G. (2019). Retinoic acid signaling pathways. Development 146. doi:10.1242/dev.167502.

28. Godsave, S., Dekker, E. J., Holling, T., Pannese, M., Boncinelli, E., and Durston, A. (1994). Expression patterns of Hoxb genes in the Xenopus embryo suggest roles in anteroposterior specification of the hindbrain and in dorsoventral patterning of the mesoderm. Dev. Biol. 166, 465–476. doi:10.1006/dbio.1994.1330.

29. Grandel, H., Lun, K., Rauch, G.-J., Rhinn, M., Piotrowski, T., Houart, C., Sordino, P., Küchler, A. M., Schulte-Merker, S., Geisler, R., et al. (2002). Retinoic acid signalling in the zebrafish embryo is necessary during pre-segmentation stages to pattern the anterior-posterior axis of the CNS and to induce a pectoral fin bud. Development 129, 2851–2865.

30. Hartomo, T. B., Van Huyen Pham, T., Yamamoto, N., Hirase, S., Hasegawa, D., Kosaka, Y., Matsuo, M., Hayakawa, A., Takeshima, Y., Iijima, K., et al. (2015). Involvement of aldehyde dehydrogenase 1A2 in the regulation of cancer stem cell properties in neuroblastoma. Int. J. Oncol. 46, 1089–1098. doi:10.3892/ijo.2014.2801.

31. Hollemann, T., Chen, Y., Grunz, H., and Pieler, T. (1998). Regionalized metabolic activity establishes boundaries of retinoic acid signalling. EMBO J. 17, 7361– 7372. doi:10.1093/emboj/17.24.7361.

32. Huang, D. W., Sherman, B. T., and Lempicki, R. A. (2009). Systematic and integrative analysis of large gene lists using DAVID bioinformatics resources. Nat. Protoc. 4, 44–57. doi:10.1038/nprot.2008.211.

33. Janesick, A., Abbey, R., Chung, C., Liu, S., Taketani, M., and Blumberg, B. (2013). ERF and ETV3L are retinoic acid-inducible repressors required for primary neurogenesis. Development 140, 3095–3106. doi:10.1242/dev.093716.

34. Janesick, A., Nguyen, T. T. L., Aisaki, K., Igarashi, K., Kitajima, S., Chandraratna, R. A. S., Kanno, J., and Blumberg, B. (2014). Active repression by RARγ signaling is required for vertebrate axial elongation. Development 141, 2260–2270. doi:10.1242/dev.103705.

35. Janesick, A., Wu, S. C., and Blumberg, B. (2015). Retinoic acid signaling and neuronal differentiation. Cell. Mol. Life Sci. 72, 1559–1576. doi:10.1007/s00018-014-1815-9.

36. Johnson, W. E., Li, C., and Rabinovic, A. (2007). Adjusting batch effects in microarray expression data using empirical Bayes methods. Biostatistics 8, 118–127. doi:10.1093/biostatistics/kxj037.

37. Kam, R. K. T., Shi, W., Chan, S. O., Chen, Y., Xu, G., Lau, C. B.-S., Fung, K. P., Chan, W. Y., and Zhao, H. (2013). Dhrs3 protein attenuates retinoic acid signaling and is required for early embryonic patterning. J. Biol. Chem. 288, 31477–31487. doi:10.1074/jbc.M113.514984.

38. Karimi, K., Fortriede, J. D., Lotay, V. S., Burns, K. A., Wang, D. Z., Fisher, M. E., Pells, T. J., James-Zorn, C., Wang, Y., Ponferrada, V. G., et al. (2018). Xenbase: a genomic, epigenomic and transcriptomic model organism database. Nucleic Acids Res. 46, D861–D868. doi:10.1093/nar/gkx936.

39. Kedishvili, N. Y. (2013). Enzymology of retinoic acid biosynthesis and degradation. J. Lipid Res. 54, 1744–1760. doi:10.1194/jlr.R037028.

40. Kedishvili, N. Y. (2016). Retinoic acid synthesis and degradation. Subcell Biochem 81, 127–161. doi:10.1007/978-94-024-0945-1_5.

41. Kessel, M., and Gruss, P. (1991). Homeotic transformations of murine vertebrae and concomitant alteration of Hox codes induced by retinoic acid. Cell 67, 89–104. doi:10.1016/0092-8674(91)90574-i.

42. Kessel, M. (1992). Respecification of vertebral identities by retinoic acid. Development 115, 487–501.

43. Kim, H., Lapointe, J., Kaygusuz, G., Ong, D. E., Li, C., van de Rijn, M., Brooks, J. D., and Pollack, J. R. (2005). The retinoic acid synthesis gene ALDH1a2 is a candidate tumor suppressor in prostate cancer. Cancer Res. 65, 8118–8124. doi:10.1158/0008-5472.CAN-04-4562.

44. Koide, T., Downes, M., Chandraratna, R. A., Blumberg, B., and Umesono, K. (2001). Active repression of RAR signaling is required for head formation. Genes Dev. 15, 2111–2121. doi:10.1101/gad.908801.

45. Kot-Leibovich, H., and Fainsod, A. (2009). Ethanol induces embryonic malformations by competing for retinaldehyde dehydrogenase activity during vertebrate gastrulation. Dis. Model. Mech. 2, 295–305. doi:10.1242/dmm.001420.

46. Kraft, J. C., Kimelman, D., and Juchau, M. R. (1995). Xenopus laevis: a model system for the study of embryonic retinoid metabolism. I. Embryonic metabolism of 9-cis- and all-trans-retinals and retinols to their corresponding acid forms. Drug Metab. Dispos. 23, 72–82.

47. Kraft, J. C., Schuh, T., Juchau, M., and Kimelman, D. (1994). The retinoid X receptor ligand, 9-cis-retinoic acid, is a potential regulator of early Xenopus development. Proc Natl Acad Sci USA 91, 3067–3071.

48. Kuttippurathu, L., Juskeviciute, E., Dippold, R. P., Hoek, J. B., and Vadigepalli, R. (2016). A novel comparative pattern analysis approach identifies chronic alcohol mediated dysregulation of transcriptomic dynamics during liver regeneration. BMC Genomics 17, 260. doi:10.1186/s12864-016-2492-x.

49. Langfelder, P., and Horvath, S. (2008). WGCNA: an R package for weighted correlation network analysis. BMC Bioinformatics 9, 559. doi:10.1186/1471-2105-9-559.

50. Lee, L. M. Y., Leung, C.-Y., Tang, W. W. C., Choi, H.-L., Leung, Y.-C., McCaffery, P. J., Wang, C.-C., Woolf, A. S., and Shum, A. S. W. (2012). A paradoxical teratogenic mechanism for retinoic acid. Proc Natl Acad Sci USA 109, 13668– 13673. doi:10.1073/pnas.1200872109.

51. Livak, K. J., and Schmittgen, T. D. (2001). Analysis of relative gene expression data using real-time quantitative PCR and the 2(-Delta Delta C(T)) Method. Methods 25, 402–408. doi:10.1006/meth.2001.1262.

52. Lohnes, D., Mark, M., Mendelsohn, C., Dollé, P., Decimo, D., LeMeur, M., Dierich, A., Gorry, P., and Chambon, P. (1995). Developmental roles of the retinoic acid receptors. J. Steroid Biochem. Mol. Biol. 53, 475–486.

53. Love, M. I., Huber, W., and Anders, S. (2014). Moderated estimation of fold change and dispersion for RNA-seq data with DESeq2. Genome Biol. 15, 550. doi:10.1186/s13059-014-0550-8.

54. Lupo, G., Liu, Y., Qiu, R., Chandraratna, R. A. S., Barsacchi, G., He, R.-Q., and Harris, W. A. (2005). Dorsoventral patterning of the Xenopus eye: a collaboration of Retinoid, Hedgehog and FGF receptor signaling. Development 132, 1737–1748. doi:10.1242/dev.01726.

55. le Maire, A., and Bourguet, W. (2014). Retinoic acid receptors: structural basis for coregulator interaction and exchange. Subcell Biochem 70, 37–54. doi:10.1007/978-94-017-9050-5_3.

56. Marshall, H., Nonchev, S., Sham, M. H., Muchamore, I., Lumsden, A., and Krumlauf, R. (1992). Retinoic acid alters hindbrain Hox code and induces transformation of rhombomeres 2/3 into a 4/5 identity. Nature 360, 737–741. doi:10.1038/360737a0.

57. Metzler, M. A., and Sandell, L. L. (2016). Enzymatic metabolism of vitamin A in developing vertebrate embryos. Nutrients 8, pii: E812. doi:10.3390/nu8120812.

58. Miettinen, K. (1998). *Nonlinear Multiobjective Optimization*. Boston, MA: Springer US doi:10.1007/978-1-4615-5563-6.

59. Moss, J. B., Xavier-Neto, J., Shapiro, M. D., Nayeem, S. M., McCaffery, P., Dräger, U. C., and Rosenthal, N. (1998). Dynamic patterns of retinoic acid synthesis and response in the developing mammalian heart. Dev. Biol. 199, 55–71. doi:10.1006/dbio.1998.8911.

60. Napoli, J. L. (2017). Cellular retinoid binding-proteins, CRBP, CRABP, FABP5: Effects on retinoid metabolism, function and related diseases. Pharmacol. Ther. 173, 19–33. doi:10.1016/j.pharmthera.2017.01.004.

61. Niederreither, K., McCaffery, P., Dräger, U. C., Chambon, P., and Dollé, P. (1997). Restricted expression and retinoic acid-induced downregulation of the retinaldehyde dehydrogenase type 2 (RALDH-2) gene during mouse development. Mech. Dev. 62, 67–78. doi:10.1016/S0925-4773(96)00653-3.

62. Niederreither, K., Subbarayan, V., Dollé, P., and Chambon, P. (1999). Embryonic retinoic acid synthesis is essential for early mouse post-implantation development. Nat. Genet. 21, 444–448. doi:10.1038/7788.

63. Nieuwkoop, P. D., and Faber, J. (1967). Normal table of Xenopus laevis (Daudin): A systematical and chronological survey of the development from the fertilized egg till the end of metamorphosis. Amsterdam: North-Holland Publishing Company.

64. Nijhout, H. F., Best, J. A., and Reed, M. C. (2019). Systems biology of robustness and homeostatic mechanisms. Wiley Interdiscip. Rev. Syst. Biol. Med. 11, e1440. doi:10.1002/wsbm.1440.

65. Paganelli, A., Gnazzo, V., Acosta, H., López, S. L., and Carrasco, A. E. (2010). Glyphosate-based herbicides produce teratogenic effects on vertebrates by impairing retinoic acid signaling. Chem. Res. Toxicol. 23, 1586–1595. doi:10.1021/tx1001749.

66. Pangilinan, F., Molloy, A. M., Mills, J. L., Troendle, J. F., Parle-McDermott, A., Kay, D. M., Browne, M. L., McGrath, E. C., Abaan, H. O., Sutton, M., et al. (2014). Replication and exploratory analysis of 24 candidate risk polymorphisms for neural tube defects. BMC Med. Genet. 15, 102. doi:10.1186/s12881-014-0102-9.

67. Papalopulu, N., Clarke, J. D., Bradley, L., Wilkinson, D., Krumlauf, R., and Holder, N. (1991). Retinoic acid causes abnormal development and segmental patterning of the anterior hindbrain in Xenopus embryos. Development 113, 1145–1158.

68. Pavez Loriè, E., Chamcheu, J. C., Vahlquist, A., and Törmä, H. (2009). Both all-trans retinoic acid and cytochrome P450 (CYP26) inhibitors affect the expression of vitamin A metabolizing enzymes and retinoid biomarkers in organotypic epidermis. Arch. Dermatol. Res. 301, 475–485. doi:10.1007/s00403-009-0937-7.

69. Porté, S., Xavier Ruiz, F., Giménez, J., Molist, I., Alvarez, S., Domínguez, M., Alvarez, R., de Lera, A. R., Parés, X., and Farrés, J. (2013). Aldo-keto reductases in retinoid metabolism: search for substrate specificity and inhibitor selectivity. Chem. Biol. Interact. 202, 186–194. doi:10.1016/j.cbi.2012.11.014.

70. Reijntjes, S., Blentic, A., Gale, E., and Maden, M. (2005). The control of morphogen signalling: regulation of the synthesis and catabolism of retinoic acid in the developing embryo. Dev. Biol. 285, 224–237. doi:10.1016/j.ydbio.2005.06.019.

71. Reijntjes, S., Zile, M. H., and Maden, M. (2010). The expression of Stra6 and Rdh10 in the avian embryo and their contribution to the generation of retinoid signatures. Int. J. Dev. Biol. 54, 1267–1275. doi:10.1387/ijdb.093009sr.

72. Ritchie, M. E., Phipson, B., Wu, D., Hu, Y., Law, C. W., Shi, W., and Smyth, G. K. (2015). limma powers differential expression analyses for RNA-sequencing and microarray studies. Nucleic Acids Res. 43, e47. doi:10.1093/nar/gkv007.

73. Romand, R., Niederreither, K., Abu-Abed, S., Petkovich, M., Fraulob, V., Hashino, E., and Dollé, P. (2004). Complementary expression patterns of retinoid acid-synthesizing and -metabolizing enzymes in pre-natal mouse inner ear structures. Gene Expr. Patterns 4, 123–133. doi:10.1016/j.modgep.2003.09.006.

74. Rydeen, A., Voisin, N., D’Aniello, E., Ravisankar, P., Devignes, C.-S., and Waxman, J. S. (2015). Excessive feedback of Cyp26a1 promotes cell non-autonomous loss of retinoic acid signaling. Dev. Biol. 405, 47–55. doi:10.1016/j.ydbio.2015.06.008.

75. Saili, K. S., Antonijevic, T., Zurlinden, T. J., Shah, I., Deisenroth, C., and Knudsen, T. B. (2019). Molecular characterization of a toxicological tipping point during human stem cell differentiation. Reprod. Toxicol. 91, 1–13. doi:10.1016/j.reprotox.2019.10.001.

76. Sakai, Y., Meno, C., Fujii, H., Nishino, J., Shiratori, H., Saijoh, Y., Rossant, J., and Hamada, H. (2001). The retinoic acid-inactivating enzyme CYP26 is essential for establishing an uneven distribution of retinoic acid along the anterio-posterior axis within the mouse embryo. Genes Dev. 15, 213–225. doi:10.1101/gad.851501.

77. Sandell, L. L., Lynn, M. L., Inman, K. E., McDowell, W., and Trainor, P. A. (2012). RDH10 oxidation of Vitamin A is a critical control step in synthesis of retinoic acid during mouse embryogenesis. PLoS ONE 7, e30698. doi:10.1371/journal.pone.0030698.

78. Savova, V., Pearl, E. J., Boke, E., Nag, A., Adzhubei, I., Horb, M. E., and Peshkin, L. (2017). Transcriptomic insights into genetic diversity of protein-coding genes in X. laevis. Dev. Biol. 424, 181–188. doi:10.1016/j.ydbio.2017.02.019.

79. Schuetz, R., Zamboni, N., Zampieri, M., Heinemann, M., and Sauer, U. (2012). Multidimensional optimality of microbial metabolism. Science 336, 601–604. doi:10.1126/science.1216882.

80. Schuh, T. J., Hall, B. L., Kraft, J. C., Privalsky, M. L., and Kimelman, D. (1993). v-erbA and citral reduce the teratogenic effects of all-trans retinoic acid and retinol, respectively, in Xenopus embryogenesis. Development 119, 785–798.

81. See, A. W.-M., Kaiser, M. E., White, J. C., and Clagett-Dame, M. (2008). A nutritional model of late embryonic vitamin A deficiency produces defects in organogenesis at a high penetrance and reveals new roles for the vitamin in skeletal development. Dev. Biol. 316, 171–190. doi:10.1016/j.ydbio.2007.10.018.

82. Session, A. M., Uno, Y., Kwon, T., Chapman, J. A., Toyoda, A., Takahashi, S., Fukui, A., Hikosaka, A., Suzuki, A., Kondo, M., et al. (2016). Genome evolution in the allotetraploid frog Xenopus laevis. Nature 538, 336–343. doi:10.1038/nature19840.

83. Shabtai, Y., Bendelac, L., Jubran, H., Hirschberg, J., and Fainsod, A. (2018). Acetaldehyde inhibits retinoic acid biosynthesis to mediate alcohol teratogenicity. Sci. Rep. 8, 347. doi:10.1038/s41598-017-18719-7.

84. Shabtai, Y., and Fainsod, A. (2018). Competition between ethanol clearance and retinoic acid biosynthesis in the induction of fetal alcohol syndrome. Biochem. Cell Biol. 96, 148–160. doi:10.1139/bcb-2017-0132.

85. Shabtai, Y., Jubran, H., Nassar, T., Hirschberg, J., and Fainsod, A. (2016). Kinetic characterization and regulation of the human retinaldehyde dehydrogenase 2 enzyme during production of retinoic acid. Biochem. J. 473, 1423–1431. doi:10.1042/BCJ20160101.

86. Shabtai, Y., Shukrun, N., and Fainsod, A. (2017). ADHFe1: a novel enzyme involved in retinoic acid-dependent Hox activation. Int. J. Dev. Biol. 61, 303–310. doi:10.1387/ijdb.160252af.

87. Sharpe, C. R., and Goldstone, K. (1997). Retinoid receptors promote primary neurogenesis in Xenopus. Development 124, 515–523.

88. Shoval, O., Sheftel, H., Shinar, G., Hart, Y., Ramote, O., Mayo, A., Dekel, E., Kavanagh, K., and Alon, U. (2012). Evolutionary trade-offs, Pareto optimality, and the geometry of phenotype space. Science 336, 1157–1160. doi:10.1126/science.1217405.

89. Sive, H. L., Draper, B. W., Harland, R. M., and Weintraub, H. (1990). Identification of a retinoic acid-sensitive period during primary axis formation in Xenopus laevis. Genes Dev. 4, 932–942. doi:10.1101/gad.4.6.932.

90. Sonneveld, E., van den Brink, C. E., van der Leede, B. M., Schulkes, R. K., Petkovich, M., van der Burg, B., and van der Saag, P. T. (1998). Human retinoic acid (RA) 4-hydroxylase (CYP26) is highly specific for all-trans-RA and can be induced through RA receptors in human breast and colon carcinoma cells. Cell Growth Differ. 9, 629–637.

91. Strate, I., Min, T. H., Iliev, D., and Pera, E. M. (2009). Retinol dehydrogenase 10 is a feedback regulator of retinoic acid signalling during axis formation and patterning of the central nervous system. Development 136, 461–472. doi:10.1242/dev.024901.

92. Taira, M., Otani, H., Jamrich, M., and Dawid, I. B. (1994). Expression of the LIM class homeobox gene Xlim-1 in pronephros and CNS cell lineages of Xenopus embryos is affected by retinoic acid and exogastrulation. Development 120, 1525–1536.

93. Tendler, A., Mayo, A., and Alon, U. (2015). Evolutionary tradeoffs, Pareto optimality and the morphology of ammonite shells. BMC Syst. Biol. 9, 12. doi:10.1186/s12918-015-0149-z.

94. Topletz, A. R., Tripathy, S., Foti, R. S., Shimshoni, J. A., Nelson, W. L., and Isoherranen, N. (2015). Induction of CYP26A1 by metabolites of retinoic acid: evidence that CYP26A1 is an important enzyme in the elimination of active retinoids. Mol. Pharmacol. 87, 430–441. doi:10.1124/mol.114.096784.

95. Urbizu, A., Toma, C., Poca, M. A., Sahuquillo, J., Cuenca-León, E., Cormand, B., and Macaya, A. (2013). Chiari malformation type I: a case-control association study of 58 developmental genes. PLoS ONE 8, e57241. doi:10.1371/journal.pone.0057241.

96. Yanai, I., Peshkin, L., Jorgensen, P., and Kirschner, M. W. (2011). Mapping gene expression in two Xenopus species: evolutionary constraints and developmental flexibility. Dev. Cell 20, 483–496. doi:10.1016/j.devcel.2011.03.015.

97. Yelin, R., Schyr, R. B.-H., Kot, H., Zins, S., Frumkin, A., Pillemer, G., and Fainsod, A. (2005). Ethanol exposure affects gene expression in the embryonic organizer and reduces retinoic acid levels. Dev. Biol. 279, 193–204. doi:10.1016/j.ydbio.2004.12.014.

98. Yu, G., Wang, L.-G., Han, Y., and He, Q.-Y. (2012). clusterProfiler: an R package for comparing biological themes among gene clusters. OMICS 16, 284–287. doi:10.1089/omi.2011.0118.

